# An integrated platform for genome engineering and gene expression perturbation in *Plasmodium falciparum*

**DOI:** 10.1101/816504

**Authors:** Armiyaw S. Nasamu, Alejandra Falla, Charisse Flerida A. Pasaje, Bridget A. Wall, Jeffrey C. Wagner, Suresh M. Ganesan, Stephen J. Goldfless, Jacquin C. Niles

**Affiliations:** Department of Biological Engineering, Massachusetts Institute of Technology, Cambridge MA 02139

## Abstract

Establishing robust genome engineering methods in the malarial parasite, *Plasmodium falciparum*, has the potential to substantially improve the efficiency with which we gain understanding of this pathogen’s biology to propel treatment and elimination efforts. Methods for manipulating gene expression and engineering the *P. falciparum* genome have been validated. However, a significant barrier to fully leveraging these advances is the difficulty associated with assembling the extremely high AT content DNA constructs required for modifying the *P. falciparum* genome. These are frequently unstable in commonly-used circular plasmids. We address this bottleneck by devising a DNA assembly framework leveraging the improved reliability with which large AT-rich regions can be efficiently manipulated in linear plasmids. This framework integrates several key functional genetics outcomes via CRISPR/Cas9 and other methods from a common, validated framework. Overall, this molecular toolkit enables *P. falciparum* genetics broadly and facilitates deeper interrogation of parasite genes involved in diverse biological processes.

## INTRODUCTION

Malaria is an infectious disease caused by five different species of *Plasmodium* parasites and continues to be a leading cause of morbidity and mortality worldwide. In 2017, there were over 219 million malaria cases and 435,000 deaths due mostly to *Plasmodium falciparum* infections (WHO, 2017). Basic molecular understanding of these parasites is still limited, and this impedes efforts to discover new therapeutic approaches to preventing, treating and eliminating malaria. The genomes of several *Plasmodium* species have now been sequenced, and nearly half of the identified genes are of uncharacterized function (Gardner et al., 2002). Therefore, functional genetics studies will be vital for systematically defining the roles of the various parasite genes toward improving our overall understanding of *Plasmodium* parasite biology. A key requirement for maximizing the effectiveness of this approach is the ability to selectively perturb gene expression to study biological function in detail.

Several recently-completed genome-scale studies have provided novel and substantial insights towards categorizing genes that are likely essential versus dispensable to survival of *Plasmodium spp* (Bushell et al., 2017; Zhang et al., 2018), and the related Apicomplexan parasite, *Toxoplasma gondii* (Sidik et al., 2016). The first comprehensive dataset came from a genome-wide CRISPR/Cas9 deletion screen in the genetically-tractable *T. gondii*. Homologous recombination-mediated knockout screens in the model rodent malaria parasite *Plasmodium berghei* (Bushell et al., 2017) and piggyBac saturation insertional mutagenesis screens in *P. falciparum* (Zhang et al., 2018) have also provided unprecedented insight into the complement of likely essential genes during asexual blood stage development in these comparably less genetically-tractable organisms. The above screening approaches facilitate identification of likely essential genes through the inability to recover surviving parasites mutated or deleted in a given locus. As such, stable cell lines with disrupted essential gene loci are usually not recovered, and this precludes direct follow-up to elucidate function. Therefore, validating the findings of these high throughput screens, and achieving detailed understanding of biological function requires implementing scalable approaches that harness methods for conditionally perturbing the expression of a rapidly expanding number of identified essential genes. Several factors impede this process. First, the most robust and reliabe conditional gene expression systems used currently in *Plasmodium* parasites require genetically encoding information at native chromosomal loci using homology-based donor vectors (Collins et al., 2013; Ganesan et al., 2016; Goldfless et al., 2014; Jones et al., 2016; Prommana et al., 2013). Second, a genomic AT content exceeding 80% (Gardner et al., 2002) complicates efficient, scalable assembly of the DNA vectors needed to achieve these modifications. Third, relatively inefficient spontaneous single- and double-crossover events have typically been used to modify target loci in a protracted process (Carvalho and Ménard, 2005). Combined, these factors complicate efforts to genetically modify many gene loci in parallel. Successful genome editing in *P. falciparum* using zinc finger nucleases (ZFNs) (Straimer et al., 2012) and CRISPR/Cas9 (Ghorbal et al., 2014; Wagner et al., 2014) provide attractive solutions for improving the ease with which targeted gene locus modification can be achieved. However, scalable and efficient application of these approaches will require facile and reliable construction of AT-rich, homology-based vectors suitable for delivering the desired final functional modifications.

Here we demonstrate a highly adaptable framework that flexibly provides reliable and scalable assembly of *P. falciparum* vectors to support stable episomal maintenance of bacterial artificial chromosomes (BACs), genome modification by single- and double-crossover integration and *Bxb1*-mediated integration (Nkrumah et al., 2006) and gene-specific editing using CRISPR/Cas9 strategies. To complement CRISPR/Cas9 genome editing experiments, we have generated a parasite cell line stably expressing the Cas9 editing machinery, which serves as a convenient alternative to co-delivery of Cas9 during transfection. We include within this framework the abilities to flexibly epitope tag, conditionally regulate, knockout and/or complement/overexpress target genes. We show that assembly of large, AT-rich regions typically encountered when creating *P. falciparum* expression and homology-directed repair vectors is readily achieved. The vectors produced using this approach successfully yield transgenic parasites in which genetic elements are either episomally maintained or chromosomally integrated as pre-specified, and that pre-installed regulatory components function as expected. Altogether, this harmonized functional genetics toolbox represents a well-validated and standardized resource that will be useful for meeting the growing and diverse needs of *P. falciparum* functional genetics.

## RESULTS and DISCUSSION

### Rationale motivating vector assembly framework

Several challenges associated with vector assembly limit the ease and scale of target-specific functional genetics studies in *P. falciparum* (Wagner et al., 2013). We defined key limitations to overcome and useful enabling functionalities to capture in our designs. We identified the need for a robust cloning chassis that supports easy manipulation and stable maintenance of AT-rich *P. falciparum* genomic DNA of broad size ranges. This permits unconstrained user selection of DNA fragment sizes best suited to the downstream application. When AT-rich sequences are present in the typical circular plasmids used to generate transgenic *P. falciparum*, they are often and unpredictably deleted and/or induce rearrangements during propagation in *E. coli* (Godiska et al., 2010). To overcome this, we selected the linear pJAZZ plasmid vector (Lucigen) as the chassis for all routine DNA assembly operations, as it has been used to successfully manipulate large AT-rich genomic fragments, including those derived from the related model rodent malaria parasite, *P. berghei* (Godiska et al., 2010; Pfander et al., 2011). We reasoned that this context would allow rapid and modular DNA assembly to be completed with high fidelity and improve overall vector construction efficiency.

While linear plasmids can facilitate accelerated DNA part assembly, only circular plasmids are stably replicated episomally in *P. falciparum* (Deitsch et al., 2001). Thus, to retain this option, a strategy for efficiently converting linear plasmids into circular forms while avoiding undesirable rearrangements is beneficial. Supercoiling in circular plasmids induces single-stranded regions within AT-rich sequences making these susceptible to nicking and deleterious rearrangements that reduce torsional stress (Benham, 1979). Therefore, rather than converting linear vectors into plasmids, we reasoned that rescuing linear vectors containing AT-rich sequences into larger BACs where supercoiling-induced torsional stress and plasmid instability are inherently lower would be highly effective. In other contexts, uncomplicated by highly AT-rich sequences, this approach has been used successfully (Guye et al., 2013). We anticipate that establishing similar approaches for *P. falciparum* will facilitate using standardized procedures to streamline and scale up production of vectors needed to pursue a range of functional genetics studies.

### Large DNA fragments containing multiple, AT-rich *P. falciparum* gene expression regulatory regions are readily ported to linear vectors

We examined the feasibility of rapidly transferring large, pre-existing DNA fragments into this linear plasmid format. This option permits direct transfer of existing parts to this new framework, while preserving access to existing user-preferred features. To demonstrate this, we transferred two configurations of our validated TetR-aptamer regulatory system that we also intended to hardwire into all future linear vector designs (Figure 1A). In the first case, an ∼7.5 kb fragment consisting of two head-to-head *P. falciparum* cassettes was transferred. One cassette encodes TetR, *Renilla* luciferase (RL) and Blasticidin S deaminase (*bsd*) as a multicistronic message using the viral 2A peptide (Straimer et al., 2012; Wagner et al., 2013). The other cassette encodes *Firefly* luciferase (FLuc) regulated by a single TetR aptamer in its 5’*UTR* (Goldfless et al., 2014). In the second case, an ∼9.5 kb fragment similar to that described above was migrated, except that the TetR component in the first cassette is replaced by a TetR–DOZI fusion, and an array of ten tandem TetR aptamers is included just upstream of the 3’*UTR* in the FLuc cassette (Ganesan et al., 2016).

**Figure 1.**
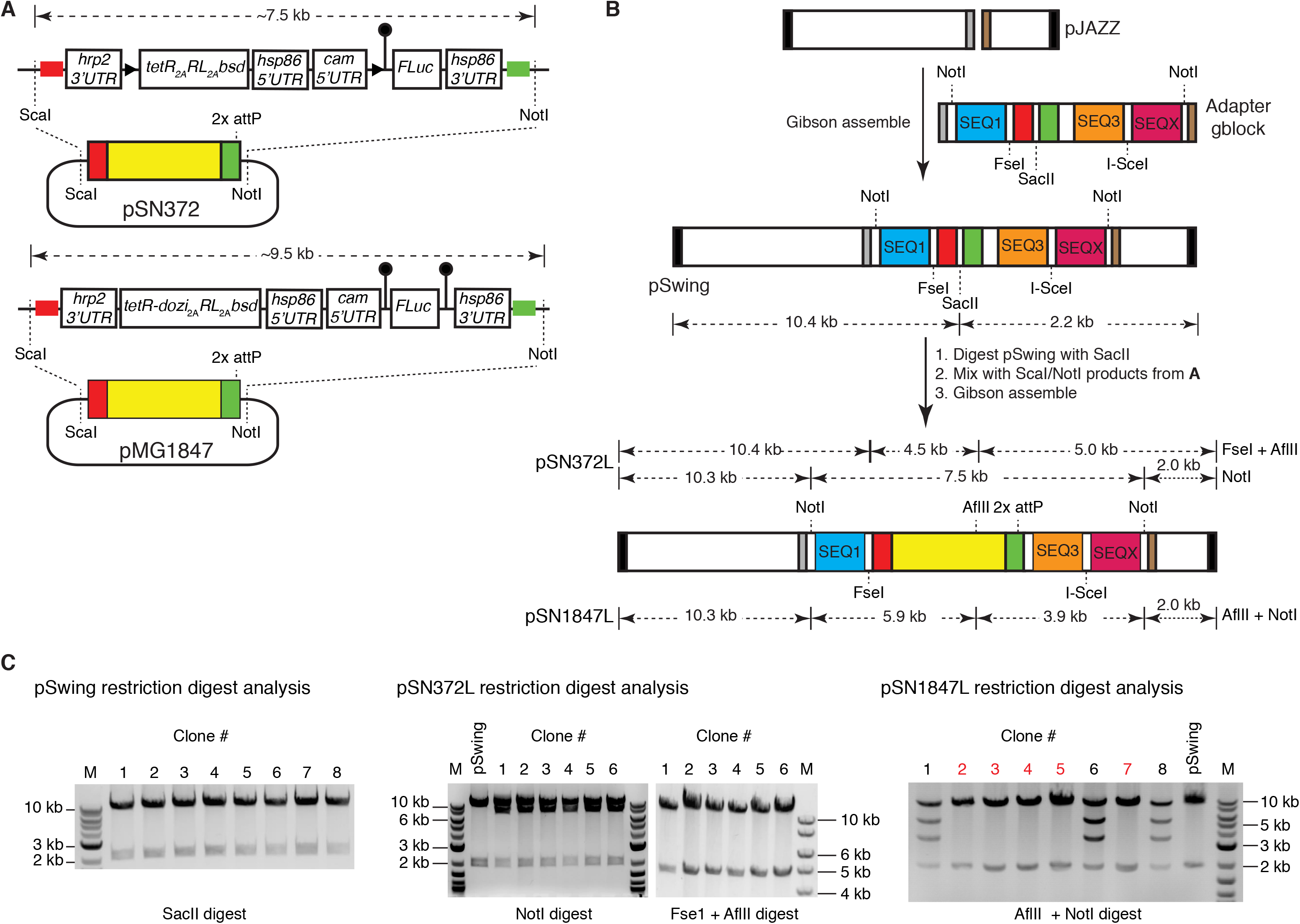
Proof-of-concept to establish successful transfer of large DNA fragments containing interspersed regions of AT-rich regulatory elements to a linear vector framework. **(A)**. The schematic shows 7.5 kb and 9.5 kb fragments to be released from extant circular vectors pSN372 and pMG1847, respectively, for assembly into linear vectors. These fragments contain TetR- or TetR-DOZI-based translation regulation modules and a transcriptional unit in which expression of a FLuc reporter CDS is translationally controlled by TetR aptamers located in either the 5’-*UTR* only or both 5’- and 3’-*UTR*s. **(B)** Strategy used to transfer the respective pSN372- and pMG1847-derived fragments into linear plasmids. The original pJAZZ-OC vector (Lucigen) was modified with a multi-cloning site gene block to create pSwing. To facilitate Gibson assembly, pSwing can be digested with restriction enzymes to expose regions homologous to cut pSN372- and pMG1847-derived fragments (red and green). **(C)**. Restriction digestion analysis confirming proper topological assembly of pSwing, pSN372L and pSN1847L. For pSN1847L, several plasmids that do not contain the expected insert, and likely corresponding to pSwing, are indicated in red font.

We opted for PCR-free transfer of these fragments to minimize the risk of introducing functionally deleterious mutations. We synthesized a gene block containing regions homologous to the fragments to be transferred using the Gibson assembly method (Gibson et al., 2009), and pre-installed this into the parental pJAZZ linear vector to create pSwing (Figure 1B and Supplementary Methods Figure 2). We released both target fragments from the circular pSN372 and pSN1847 by ScaI and NotI double digestion, cut pSwing using SacII and used single pot, 3-piece Gibson assemblies to produce pSN372L and pSN1847L (Figure 1B and Supplementary Methods Figure 3). We screened bacterial colonies by restriction enzyme mapping to identify plasmids having the expected digestion pattern (Figure 1C). These data show that large, pre-existing fragments containing AT-rich regions interspersed with regulatory components required for regulating *P. falciparum* gene expression are readily imported into this linear vector chassis.

**Figure 2.**
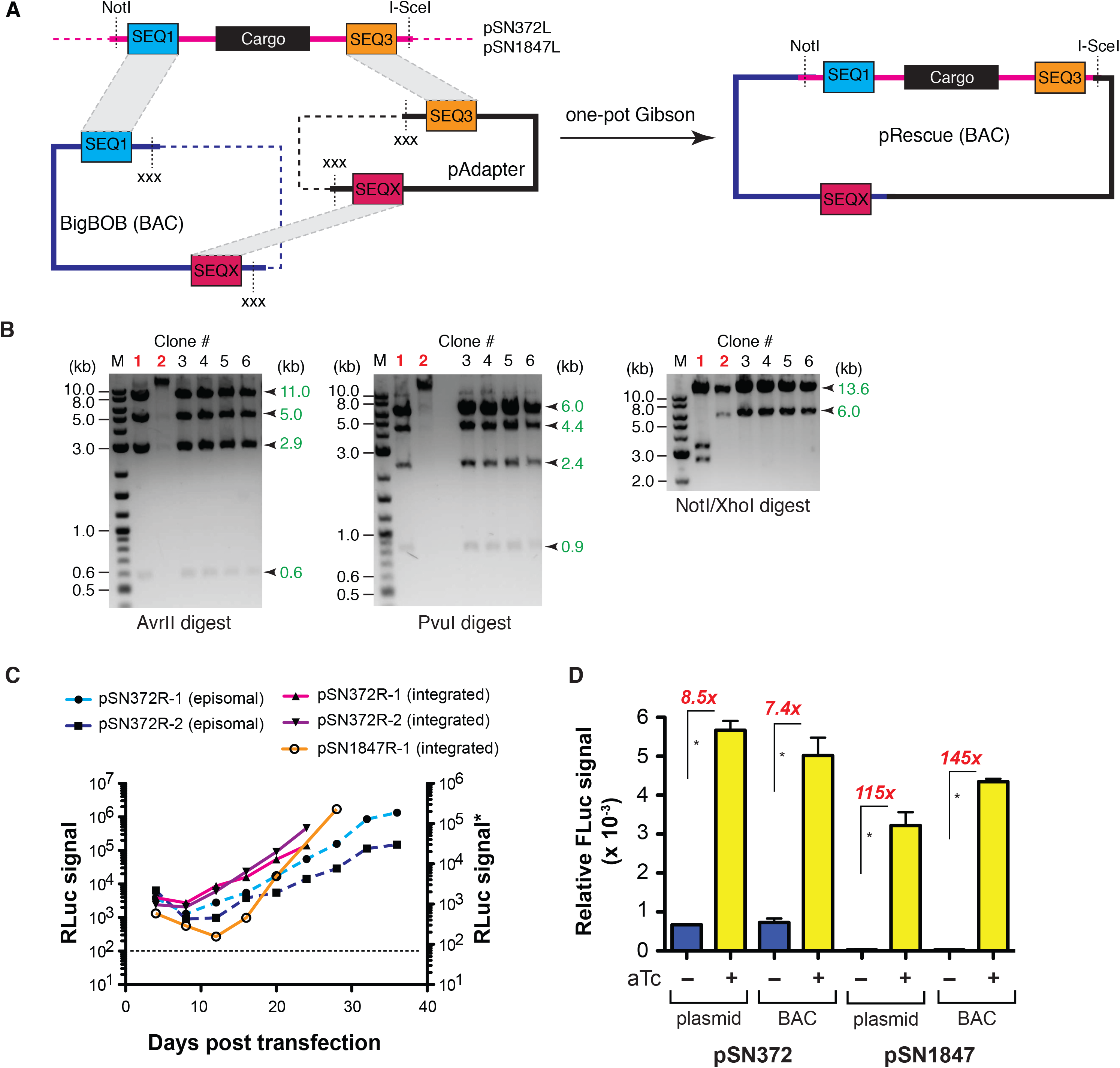
Establishing successful rescue of *P. falciparum* regulatory modules assembled in linear vectors to BACs. (**A**) Schematic of the linear vector rescue process, wherein the linear vector is digested with NotI/I-Sce1 to expose the SEQ1 and SEQX regions at the ends. The recipient BAC, BigBOB, and pAdapter plasmid are pre-configured such that the SEQ1/SEQ3 and SEQ3/SEQX regions can be exposed upon restriction enzyme digestion with PacI and XbaI/XhoI, respectively. The appropriate fragments from the linear plasmid, BigBOB BAC and pAdapter restriction enzyme digestions are mixed and assembled in a single pot, 3-piece Gibson reaction. (**B**) Restriction enzyme digestion mapping, illustrated for pSN372R, is used to identify BACs with the expected topology. Examples of BACs isolated from transformation of the same Gibson assembly reaction that were correctly (black font) versus incorrectly (red font) assembled are shown. (**C**) Renilla luciferase measurements were monitored to successful transfection of *P. falciparum* using rescued BACs. \**Note*: Renilla luciferase measurements shown in solid and dashed lines were made with the Promega Dual-Luciferase and Renilla luciferase Assay Systems, respectively; (**D**) Comparison of conditional regulation of normalized FLuc expression from plasmid and BAC contexts for pSN372 and pSN1847. Data are the mean of *n = 3* ± standard deviation for each condition. * *p* ≤ 0.05 by Student’s t-test.

**Figure 3.**
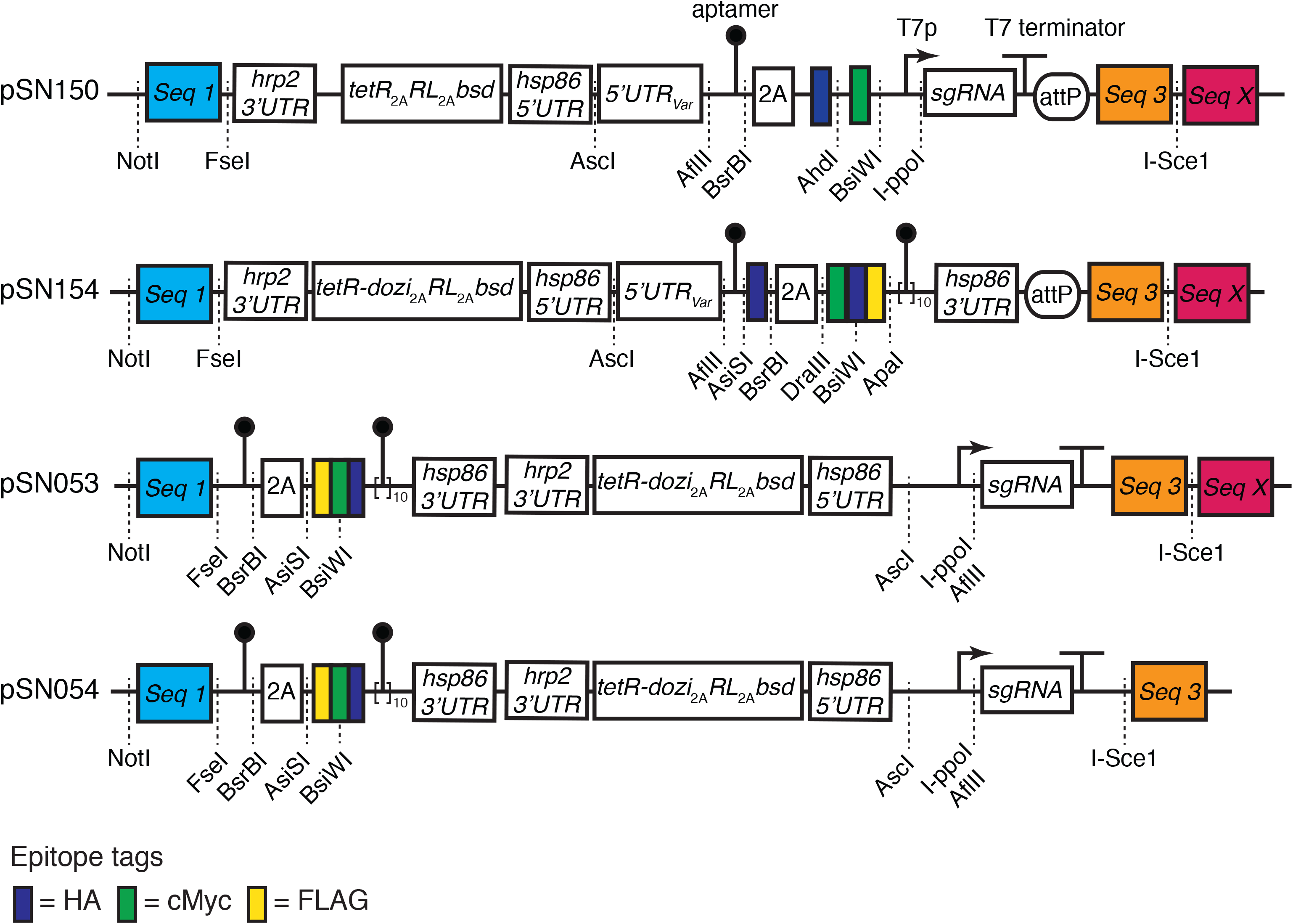
Overview of key base vectors pSN150, pSN154, pSN053 and pSN054. These can be used for enabling various modifications to native *P. falciparum* gene loci using integrase or CRISPR/Cas9 genome engineering methods described in detail in the main text.

### Linear vectors can be rescued into BACs that successfully transform *P. falciparum*

The linear vectors generated above must be converted into circular plasmids for stable episomal propagation or *Bxb1*-catalyzed chromosomal integration in *P. falciparum*. We rescued pSN372L and pSN1847L into BACs using an adapter plasmid (pAdapter) and a BAC recipient (BigBOB) as described (Guye et al., 2013). Fragments encoding components to be expressed in *P. falciparum* were released from pSN372L or pSN1847L by NotI/I-Sce1 restriction enzyme digestion, and assembled with pAdapter and BigBOB in single pot, 3-piece Gibson assembly reactions (Figure 2A). We analyzed colonies containing the rescued pSN372R and pSN1847R BACs to confirm successful transfer of the pSN372L and pSN1847L fragments, respectively. Restriction digestion mapping showed that BACs with the expected topology were obtained (Figure 2B). These data indicate that the linear vector-encoded sequence information critical for *P. falciparum* genetics studies can be readily rescued into BACs without the rearrangements frequently observed during construction of circular plasmids typically used for cloning AT-rich *P. falciparum* sequences.

### Regulatory components for controlling gene expression function predictably in BACs

Next, we sought to understand whether pSN372R and pSN1847R would: (1) successfully transform *P. falciparum*, both in episomal and chromosomally-integrated contexts; and (2) yield functional outcomes similar to those obtained using traditional *P. falciparum* expression vectors. We transfected pSN372R and pSN1847R alone for episomal maintenance, or co-transfected with a plasmid encoding the *Mycobacterium Bxb1* integrase (pINT) (Nkrumah et al., 2006) to achieve site-specific integration at the *cg6* locus in an NF54^attB^ line (Adjalley et al., 2011). We selected parasites using Blasticidin S and monitored *Renilla* luciferase signal over the transfection. *Renilla* luciferase signal increased with the expected kinetics (Wagner et al., 2013) for transfections expected to result in episomal BAC maintenance or *Bxb1*-mediated site-specific integration (Nkrumah et al., 2006) of the BAC at the *cg6* locus (Figure 2C). Taken together, these data indicate that BACs generated through this process successfully transform *P. falciparum* and the cassettes mediating *Renilla* luciferase and Blasticidin S deaminase transgene expression remain functionally intact.

We also wished to ensure that regulated gene expression mediated by the TetR and TetR–DOZI regulatory components hardwired into our linear vector and BAC frameworks was not compromised. Therefore, we measured anhydrotetracyline (aTc)-regulated expression of the *firefly* luciferase reporter gene encoded by these constructs in parasites harboring chromosomally integrated pSN372R and pSN1847R. These data show that aTc regulates *firefly* luciferase expression by 8-fold (pSN372) and 115-fold (pSN1847) when a regular plasmid is integrated, and ∼7-fold (pSN372R) and 145-fold (pSN1847R) when the rescued BACs are integrated (Figure 2D). This degree of regulation is consistent with that observed in the context of the plasmids traditionally used in *P. falciparum* (Ganesan et al., 2016; Goldfless et al., 2014). Importantly, these data demonstrate that information needed to achieve gene expression and regulation outcomes in *P. falciparum* can be encoded on BAC constructs without compromising how these DNA parts function in the parasite.

### Modular linear vectors enable easily programmable and multi-functional designs

We generated several customizable base vectors pre-configured to achieve several key outcomes desired for functional genetics studies in *P. falciparum* (Figure 3 and Supplementary Figure 1). We defined standardized architectures that collectively facilitate: promoter/5’-*UTR* swapping; N- and/or C-terminal epitope tagging; conditional regulation of gene expression; and genetic complementation by episomal or site-specific chromosomal integration. The components required for TetR aptamer-mediated regulation are hardwired into these base constructs. All variable parts needed to achieve locus-specific modification or transgene expression are modular, and we have enabled site-specific integration of any designed construct into the genome either through *Bxb1-*mediated *attB* × *attP* recombination, spontaneous single- and/or double-crossover integration or CRISPR/Cas9- and ZFN-mediated genome editing methods.

### Genetic complementation and transgene expression

pSN154 is designed to enable several outcomes, including gene complementation by either stable episome maintenance or *Bxb1* integrase-mediated chromosomal integration of rescued BACs via *attP* sites. Unique restriction sites (AsiSI, BsrBI and DraIII) modularize a region encoding a preinstalled T2A ‘skip peptide’ and HA epitope tag to allow synthesis of well-defined N-termini to either accommodate inclusion of leader peptides directing organelle-specific trafficking or optional epitope tags for protein detection. Similarly, C-terminal epitope tags (FLAG, c-Myc, HA) are preinstalled, and can be individually selected for inclusion in the encoded transgene via DraIII/BsiWI restriction sites. For regulated transgene expression, either a single and/or ten tandem TetR aptamers are preinstalled in the 5’- and 3’-*UTR*s, respectively. The 5’-UTR/promoter is modular (AscI/AflII) to allow straightforward exchange for perturbing transgene expression timing and levels (Bozdech et al., 2003b; Le Roch et al., 2003). A multi-cistronic regulatory module containing a TetR–DOZI*_T2A_*RLuc*_T2A_*Blastisidin S deaminase cassette using the T2A “skip” peptide is encoded on the same plasmid to permit regulated transgene expression (TetR–DOZI), quantitative monitoring of transfected parasites (RLuc), and positive selection of transformed parasites (Blasticidin S deaminase). This feature also ensures that all constructs using this design are compatible with any parasite strain background. Upon transgene insertion, this linear plasmid can be rescued into a BAC, and used to generate parasite lines in which the BAC is episomally maintained or chromosomally integrated (Supplementary Figure 1A).

### Programmable donor vector contexts for genome editing

#### Design 1

pSN150 is designed to simultaneously facilitate promoter swapping and conditional regulation of target gene expression by a single TetR aptamer within a user-specified synthetic 5’-*UTR*. In this case, the regulatory module consists of a TetR*_T2A_*RLuc*_T2A_*blastisidin S deaminase cassette (Figure 3). Modification of a target chromosomal locus is achieved by inserting the left homologous region (LHR) at the FseI site, and the right homologous region (RHR) using some combination of BsrBI/ApaI, AhdI and BsiWI sites, depending on the N-terminal modification desired (Supplementary Figure 1B). A modular T2A ‘skip peptide’ and epitope tag (HA and c-Myc) region is included immediately downstream of the 5’-*UTR* aptamer. Translation most likely initiates using an ATG within the aptamer sequence, and this is expected to produce an 11 amino acid N-terminal leader peptide (Goldfless et al., 2014; 2012). We have previously shown this does not interfere with proper protein trafficking to organelles (Goldfless et al., 2014). Nevertheless, we have included the T2A feature to provide the option to force exclusion of this leader from the mature target protein. Similarly, the epitope tags can be retained or excluded from the mature protein by user choice. Overall, the modularity built into this region provides the flexibility to engineer the N-terminus of targeted proteins in a manner compatible with exploring their function irrespective of subcellular trafficking and compartmentalization. This design facilitates homology-directed repair of double strand DNA breaks induced by ZFNs or CRISPR/Cas9 at the target locus. For CRISPR/Cas9 editing, we have included a cassette for producing the required sgRNA under T7 promoter control (Wagner et al., 2014). This cassette is easily modified to target a new locus by inserting the required targeting sequence via an I-PpoI/AflII site. AflII digestion is more efficient, and the preferred option.

#### Design 2

pSN053 and pSN054 are intended to extend the possibilities for accessing native gene loci via manipulation of regions upstream, downstream and within a targeted gene. pSN053 and pSN054 are easily programmed to allow installation of regulatory aptamers within the 3’*UTR* only or both 5’- and 3’-*UTR*s to achieve aTc-dependent regulation via a TetR-DOZI-containing regulatory module. These options reflect our observation that TetR–DOZI, but not TetR, enables regulated gene expression via aptamers placed in the 3’-*UTR*, and that superior dynamic regulatory range is achieved when aptamers are dually positioned in the 5’- and 3’-*UTR*s (Ganesan et al., 2016). This configuration can be reasonably achieved while either preserving expression from the native promoter or swapping promoters, if desired (Supplementary Figure 1C). More routinely, regulatory TetR aptamers can be introduced only in the 3’-*UTR* of a target coding sequence (CDS) via the relevant combination of FseI/BsrBI/AsiSI/BsiWI restriction sites (Supplementary Figure 1D). Modular N-terminal (HA) and C-terminal (FLAG, c-Myc and HA) epitope tags are preinstalled, allowing for flexible tag selection. A T7 promoter-driven cassette for producing single guide RNAs (sgRNAs) required for CRISPR/Cas9-mediated genome editing is preinstalled, and new target binding sites can be rapidly introduced via an I-PpoI/AflII site. With pSN053, the sgRNA cassette is not integrated into the parasite’s chromosome during editing, while with pSN054, this cassette is chromosomally integrated and can be used as a barcode to uniquely identify parasite lines via a standardized PCR and either Sanger or Next Generation Sequencing methods. These linear plasmids or BAC-rescued versions can be used in Cas9-mediated genome editing applications. Rescued BACs are also suitable for 3’*UTR* modification achieved by single crossover integration.

### Increasing flexible options for genome editing in *P. falciparum*

Our donor vector designs include the option for sgRNA production to facilitate Cas9-mediated genome editing. In our original implementation of CRISPR/Cas9 editing technology in *P. falciparum*, we used the orthogonal T7 RNA polymerase (T7 RNAP) to produce sgRNAs. We co-transfected parasites with a plasmid expressing *Sp*Cas9 and the sgRNA, and a donor vector expressing T7 RNAP and containing the required homology arms to repair Cas9-induced double strand breaks (Wagner et al., 2014). We have streamlined this approach to allow simultaneous expression of *Sp*Cas9 and T7 RNAP from a single pCRISPR*^hdhfr^* plasmid. T7 RNAP is produced along with the human DHFR selection marker using a T2A ‘skip peptide’ from a single expression cassette (Figure 4A). pCRISPR*^hdhfr^* contains an *attP* site to enable *Bxb1*-mediated integration into *attB* parasite lines. We have integrated pCRISPR*^hdhfr^* into an NF54*^attB^* strain (Adjalley et al., 2011) to create a clonal cell line stably expressing *Sp*Cas9 and T7 RNAP proteins (Figure 4B), which can serve as a convenient background for the various described genome editing outcomes.

**Figure 4.**
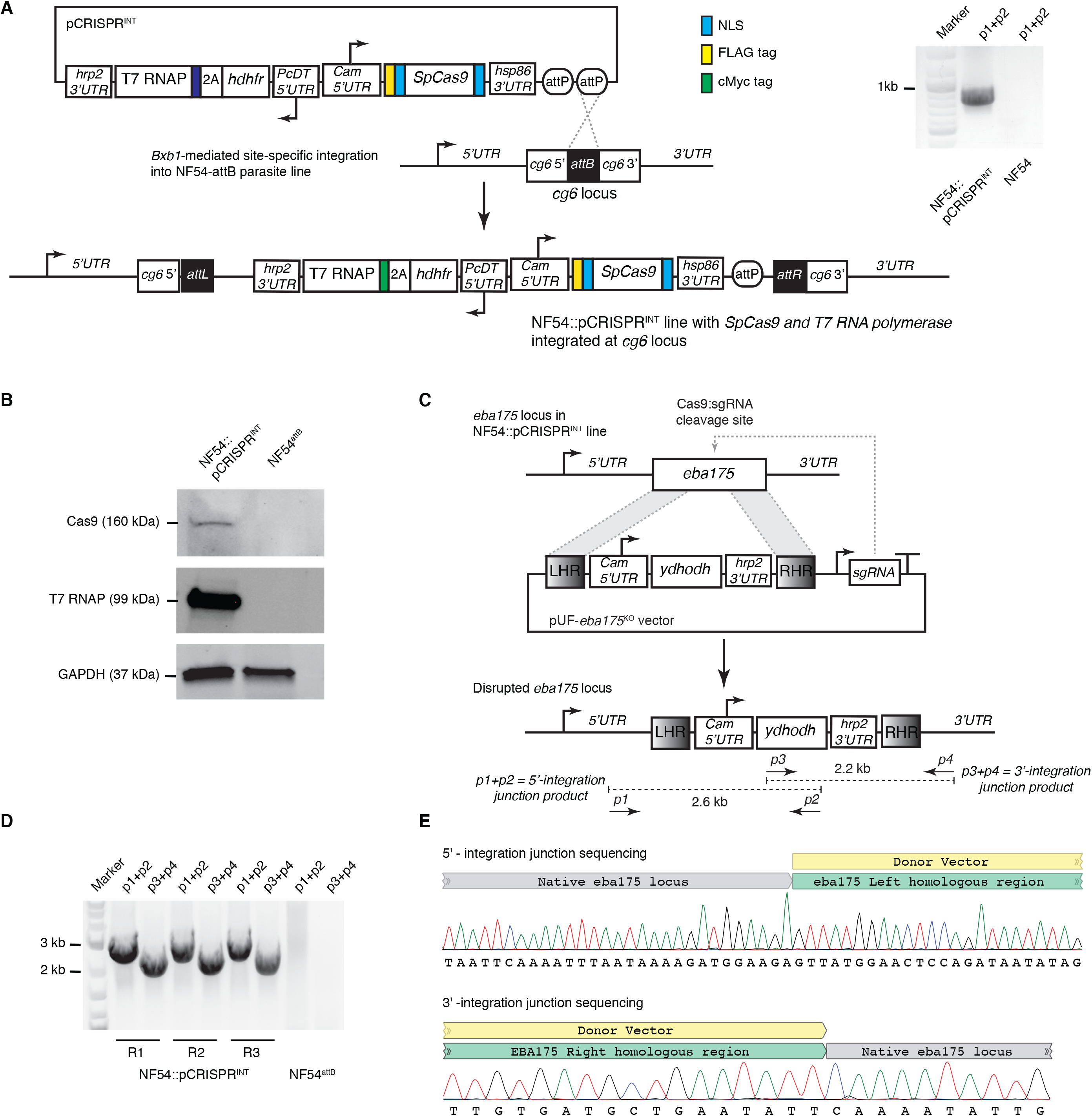
Validation of a pCRISPR plasmid reagent for CRISPR/Cas9-mediated genome engineering via transient or constitutive *Sp*Cas9 expression. (**A**) The pCRISPR plasmid allows constitutive expression of *Sp*Cas9 from the CAM 5’-*UTR*/hsp86 3’-*UTR* transcription unit. Through use of a T2A “skip peptide”, both the human dihydrofolate reductase (*hdhfr*) selectable marker and T7 RNA polymerase for sgRNA transcription are expressed from the *Pc*DT 5’-*UTR*/hrp2 3’-*UTR* transcription unit. The 2x *attP* sites allow for *Bxb1*-mediated integration into a chromosomal *attB* site previously engineered into a host parasite strain. Successful integration of pCRISPR plasmid at the *cg6* locus in an NF54::pCRISPR parasite line was confirmed by PCR analysis. (**B**) Western blot analysis to detect expression of *Sp*Cas9 (anti-FLAG) and T7 RNA polymerase (anti-Myc) proteins by the NF54::pCRISPR parasite. (**C**) Schematic of the donor vector used to disrupt the dispensable *eba175* gene, and configuration of the *eba175* locus in the NF54::pCRISPR line pre- and post-editing. The donor vector contained left and right homologous regions from *eba175* flanking a selectable *ydhodh* expression cassette, and a T7 RNAP-transcribed cassette for expression of the sgRNA targeting *eba175*. (**D**) PCR analysis to detect the 5’- and 3’-integration junctions in edited lines using the primer pairs *p1+p2* and *p3+p4*, respectively. (**E**) Sanger sequencing data for the 5’- and 3’-integration junction PCR products obtained from the knockout transfection replicate, R3.

To validate that the NF54::pCRISPR line is competent for genome editing applications, we transfected this line with a homology-directed repair vector designed to disrupt the dispensable *eba175* invasion ligand through insertion of a DSM-1 selectable *ydhodh* expression cassette (Figure 4C). We recovered viable parasites from independent triplicate transfections. In all cases, PCR and sequencing analyses confirmed that the expected insertion event into the *eba175* locus had been successfully achieved (Figure 4D,E and Supplementary Figure 2). Altogether, these new reagents increase the flexibility with which the *P. falciparum* genome can be edited either through co-transfection of donor vectors with the pCRISPR*^hdhfr^* plasmid into a user-specified parasite strain, or by transfecting donor vectors into established pCRISPR*^hdhfr^* cell lines.

### Linear plasmids facilitate donor vector assembly for engineering the *P. falciparum* genome to enable functional studies

To establish general applicability of this framework for functional genetics in *P. falciparum*, we sought next to demonstrate successful: (1) assembly of target constructs into the described linear vectors (pSN150 and pSN053/4, specifically) and successful rescue into BACs; and (2) use of these vectors to conveniently create genetically engineered parasites compatible with performing future biological studies aimed at studying parasite gene function in finer detail. We focused primarily on designs aimed at manipulating native gene loci.

### 1. Swapping native promoters for TetR aptamer-regulated synthetic promoters using CRISPR/Cas9 genome editing

The timing and level of global gene expression during blood stage malaria parasite development is tightly regulated by both transcriptional and post-transcriptional mechanisms (Bozdech et al., 2003a; Bunnik et al., 2013; Caro et al., 2014; Le Roch et al., 2003). The ability to perturb both of these gene expression parameters, therefore, can provide useful insights into how essential cell cycle, metabolic, and other biological processes are coordinated to ensure proper development. For example, previous work has shown that temporally ectopic expression of many parasite genes occurs when expression of CCR4-Associated Factor 1 (CAF1), a key regulator of mRNA metabolism, is disrupted. This resulted in severely dysregulated expression of genes involved in egress and invasion, and development of parasites in red blood cell stages was severely impaired (Balu et al., 2011).

Therefore, we sought to establish the technical feasibility of engineering several gene loci to replace native promoters with a non-cognate promoter/5’-*UTR* regulated by a single TetR aptamer. We selected five target genes for this proof-of-concept to sample both putatively essential and dispensable genes: choline kinase (CK; <u>PF3D7_1401800</u>); chloroquine resistance transporter (CRT; <u>PF3D7_0709000</u>); glycogen synthase kinase (GSK3; <u>PF3D7_0312400</u>); hexose transporter (HT; <u>PF3D7_0204700</u>); and thioredoxin reductase (TrxR; <u>PF3D7_0923800</u>). Except TrxR, these genes were considered to be likely essential in *P. falciparum* based on an insertional mutagenesis screen (Zhang et al., 2018). In the related *P. berghei*, only HT and CRT were considered likely essential, while CK, GSK3 and TrxR were dispensable (Bushell et al., 2017). These data suggest that CRT and HT are likely essential with highest confidence, and that knocking down their expression would more likely confer losses in parasite fitness and growth. Even with substantial knockdown, no functional outcome would be expected if CK, GSK3 and TrxR are dispensable in accordance with the *P. berghei* data. On the other hand, if essential as suggested by the piggyBAC screen, even modest knockdown levels could result in loss of parasite fitness.

We used a 625 bp fragment derived from the *P. falciparum* calmodulin (Cam) promoter (Crabb and Cowman, 1996) modified by a single, regulatory TetR aptamer 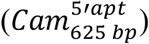 as the non-cognate, synthetic promoter/5’-*UTR* to replace the cognate promoter/5’-*UTR*s in these experiments. Relatively few instances of functional 5’-*UTR* manipulation in *Plasmodium* blood stages have been reported (Goldfless et al., 2014; Pino et al., 2012). This reflects a combination of: (1) technical difficulties associated with 5’-*UTR* engineering using previous approaches; and (2) the challenge of ensuring quantitatively adequate expression of essential proteins to allow parasite survival at baseline, and then triggering sufficient protein depletion such that levels fall below the functional threshold. Using pSN150, we designed and assembled constructs to exchange the native promoter, and install an HA tag at the expected N-terminus of each target protein (Figure 5A). Upstream and downstream homologous regions needed for editing the intended locus were PCR amplified from within the native 5’-*UTR* and target gene, respectively. To ensure that repair is irreversible and simultaneously preserve the option to use the same donor vector with alternative sgRNAs, we synthesized recoded DNA segments beginning at the ATG of the target coding sequence through a region containing several candidate Cas9 target sites.

**Figure 5.**
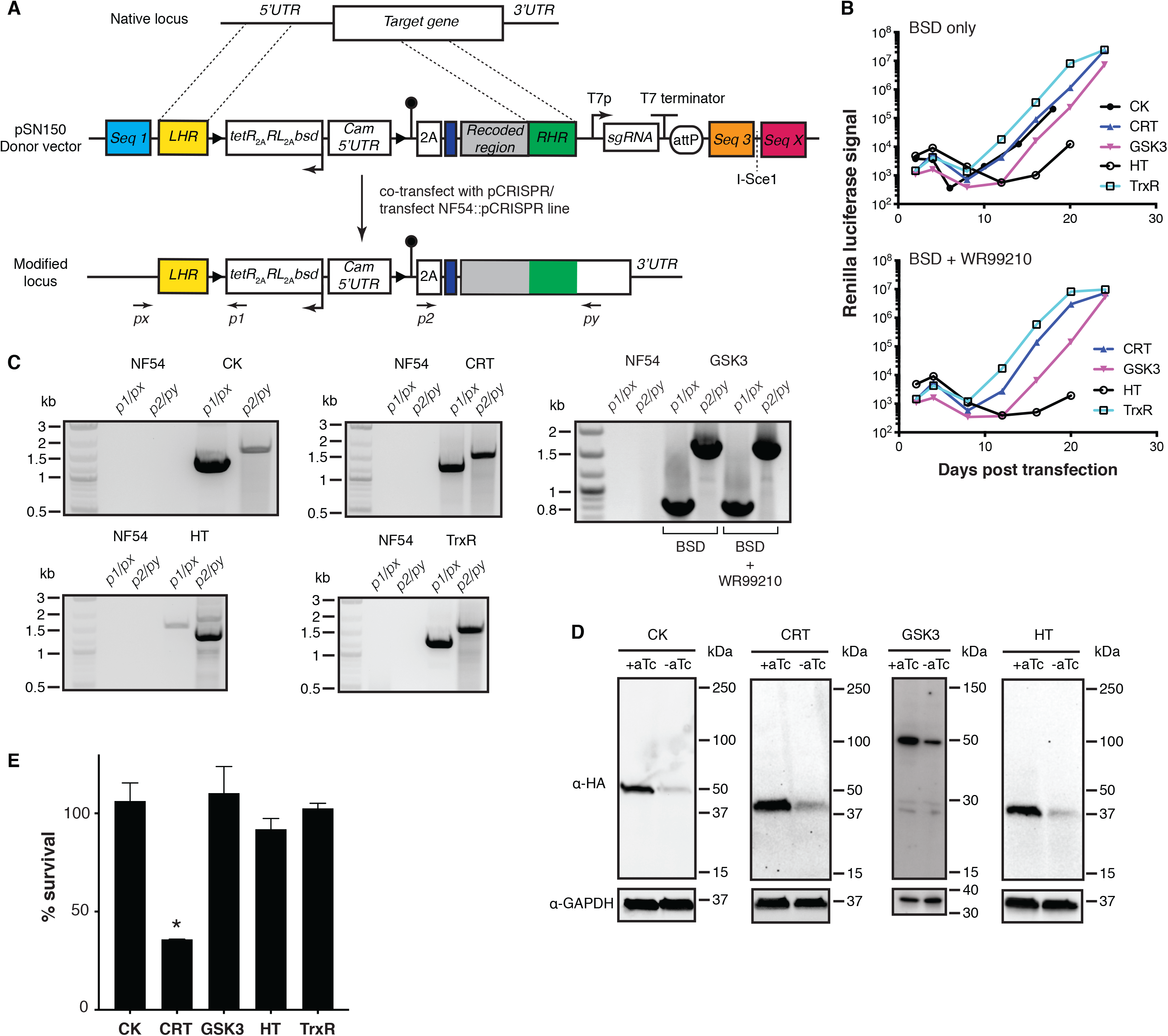
Manipulating *5’-UTRs* of native loci using BAC-rescued pSN150 donor vectors and CRISPR/Cas9 genome editing. (**A**) Generalized schematic of CRISPR/Cas9-mediated modification of a target locus to install a synthetic 5’UTR regulated by a single TetR-binding aptamer, an N-terminal HA tag and an expression cassette producing TetR, *Renilla* luciferase (RL) and the Blasticidin S deaminase selection marker. Engineered parasites were generated by either co-transfecting donor vectors with the pCRISPR plasmid or transfecting into the NF54::pCRISPR line described in Figure 4. (**B**) Successful generation of stable parasite lines was monitored via *Renilla* luciferase levels, under conditions where only the donor vector was positively selected (BSD) or both donor vector and pCRISPR plasmids were selected (BSD+WR99210). (**C**) gDNA extracted from transgenic parasites analyzed by PCR demonstrated formation of the expected *5’*- and *3’*- integration junctions at each targeted locus. Expected junctional PCR product sizes (*5’*, *3’*): *CK* (1.3 kb, 1.7 kb)*; CRT* (1.2 kb, 1.6 kb); *GSK3* (0.84 kb, 1.68 kb); *HT* (1.6 kb,1.6 kb); and *TrxR* (1.3 kb,1.7 kb). Marker = 1 kb Plus DNA ladder (New England Biolabs). (**D**) Western blot analysis of target protein expression under ± aTc conditions. Expected molecular weights of the HA-tagged proteins are: CK = 53.7 kDa; CRT = 50.2 kDa; GSK = 51.6 kDa; and HT = 57.9 kDa. *Note*: HT migrates faster than its expected molecular weight, but identically when detected by an N- and C-terminal tag (Figure 7). (**E**) Normalized *Renilla* luciferase levels or SYBR Green I staining (GSK3 line only) for parasites grown in the absence (0 nM) or presence (50 nM) aTc for 72 h. Data represent the mean of *n = 3* ± s.e.m for CK, CRT, HT, and TrxR, and *n = 2* ± s.e.m for GSK3. * *p* ≤ 0.05 by Student’s t-test.

We co-transfected NF54^attB^ parasites with pSN150 donor BACs targeting CK, CRT, GSK3, HT and TrxR with pCRISPR*^hdhfr^* in the presence of aTc to generate edited lines. We selected for either the donor BAC only (BSD) or both donor BAC and pCRISPR (BSD + WR99210), and successfully recovered transformed parasites using both selection protocols (Figure 5B). This confirmed suitability of pSN150-derived BACs for generating transgenic parasites, and adequate production of putatively essential CRT and HT proteins using a non-cognate synthetic promoter. Using PCR and sequencing, we verified that parasites were edited as expected. For all target genes, independently of whether the pCRISPR*^hdhfr^* plasmid was co-selected, we detected the expected integration events in transgenic lines, but not the parental NF45*^attB^* (Figure 5C). Western blot analysis revealed aTc-dependent regulation of CK, CRT, GSK3 and HT protein expression (Figure 5D). TrxR could not be detected, even though DNA sequencing confirmed the epitope tag was in-frame with the TrxR coding sequence. This observation is consistent with the previous studies showing that two isoforms of TrxR are produced: a cytosolic form initiated from an internal, alternate translation initiation site; and a mitochondrial form resulting from N-terminal processing of the expected full-length protein (Kehr et al., 2010). Therefore, it is likely that mature TrxR produced by the parasite excludes the N-terminal epitope tag.

We compared aTc-dependent growth of these nonclonal engineered parasites. No significant aTc-dependent difference in relative growth was observed for the engineered CK, GSK3 and TrxR lines, but growth of the engineered CRT line decreased by ∼60% under knockdown conditions (Figure 5E). For the engineered HT line grown in standard RPMI with 2 mg/mL glucose, no aTc-dependence in growth was detected. As HT mediates glucose transport, we also evaluated growth under more limiting glucose conditions. In 0.2 mg/mL glucose there was ∼50% reduction in growth in the absence of aTc (Supplementary Figure 3). These data show that, consistent with their expected essential functions, conditional knockdown of CRT and HT expression levels produced measurable reductions in parasite growth. In contrast, no such growth deficits were detected with the potentially dispensable CK, GSK3 and TrxR knockdown lines.

Given the discordance in data between *P. falciparum* and *P. berghei* with respect to CK and GSK3 essentiality, we used the pSN150 framework to generate knockouts in *P. falciparum*. Beginning with the vectors used to modify the 5’-*UTR*s of CK and GSK3, we preserved the *bsd* selection cassette, left homologous region and sgRNA cassette, but replaced the fragment containing the synthetic promoter through to the original right homologous region with a new right homologous region overlapping with the 3’-*UTR* of the targeted locus (Figure 6A). We included a similarly designed construct for TrxR as a control for a non-essential gene, as direct transposon insertion into the coding sequence at this locus was detected in the *piggyBAC* screen (Zhang et al., 2018). We transfected linear plasmids, each in duplicate, into the NF54::pCRISPR*^hdhfr^* line under BSD selection pressure, since successful homology directed repair of the target locus will result in chromosomal integration of the BSD resistance to facilitate selection. In all instances, we recovered BSD-resistant, RLuc positive parasites post-transfection. PCR and sequencing analysis of genomic DNA isolated from these parasites revealed that successful disruption of the targeted locus had been achieved in all instances (Figure 6B-D). Thus, consistent with the *P. berghei* data, our findings indicate that CK, GSK3 and TrxR are dispensable during *P. falciparum* blood stage growth.

**Figure 6.**
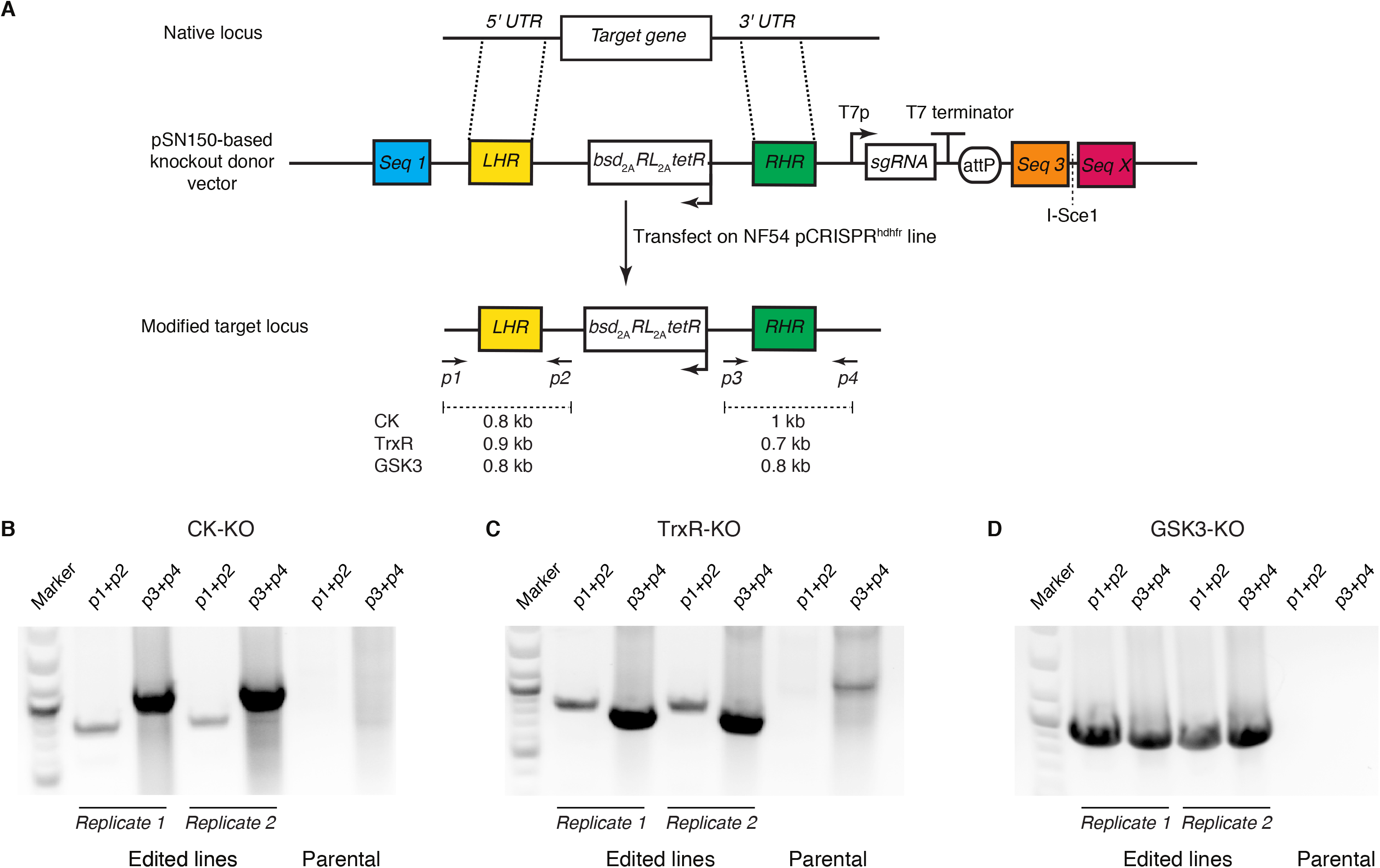
Linear pSN150 can be used for efficient CRISPR/Cas9-mediated deletion of non-essential *P. falciparum* loci. (**A**) Schematic of a pSN150 vector configured to delete a target locus via homology-directed repair after Cas9-induced cleavage mediated by a sgRNA transcribed from the same vector. (**B-D**) PCR analysis to detect the 5’- and 3’-integration junctions diagnostic of successfully edited parasite lines using the primer pairs *p1+p2* and *p3+p4*, respectively. *p1* and *p4* primers were selected to recognize the CK, TrxR and GSK3 loci being targeted, while *p2* and *p3* are common primers that bind, respectively, within the *hrp2* 3’- *UTR* and hsp86 5’-*UTR* common to all pSN150-derived knockout vectors.

Altogether, these data indicate that genome engineering using easily-designed pSN150-based donor vectors and CRISPR/Cas9 methods can be used to efficiently reconfigure promoter/5’- *UTR* regions in their native chromosomal context. Appropriately selected synthetic promoters can also be substituted for native promoters to achieve regulated and functionally informative perturbation of essential target protein expression. Furthermore, pSN150-based donor vectors can be conveniently reconfigured to create targeted gene deletion vectors that can be used in conjunction with parasite cell lines stably expressing Cas9 to evaluate gene essentiality. These easy-to-implement and high-resolution gene deletion manipulations are complementary to recently-described genome-scale approaches (Bushell et al., 2017; Sidik et al., 2016; Zhang et al., 2018) as tools to rigorously validate gene essentiality, as highlighted here for CK and GSK3. The potential to scale up production of these vectors provides opportunities for use in primary screens implemented with high gene-targeting efficiency and specificity, while introducing well-defined loss-of-function genetic changes. Such desired outcomes may not be consistently attainable in random insertional mutagenesis screens, especially if full saturation is not achieved. Altogether, the combined use of these various approaches will be crucial for most accurately characterizing essential gene function in *P. falciparum*.

### 2. Native allele modification to install C-terminal epitope tags and regulatory TetR aptamers in 3’-*UTR* and both 5’- and 3’-*UTR*s by CRISPR/Cas9 genome editing

Modifying the 3’-*UTR* of genes in their chromosomal contexts permits insertion of epitope tags and regulatory elements while preserving native timing and similar expression levels associated with the endogenous promoter. This is desirable in functional studies requiring perturbations in protein levels that more precisely match physiological expression timing so as to facilitate elucidation of biological function.

To achieve this, we have used pSN053 and/or pSN054 together with CRISPR/Cas9 genome editing to readily modify diverse loci (Figure 7A and Supplementary Figure 1C and 1D). Here, we show this by targeting the hexose transporter (HT; <u>PF3D7_0204700</u>), ferrochelatase (FC; <u>PF3D7_1364900</u>) and a putative amino acid transporter (AAT; <u>PF3D7_0209600</u>) using pSN054-based donor vectors. Once sgRNAs targeting a site near the 3’-*UTR* of these genes were selected, left homologous regions together with a recoded region overlapping the sgRNA target site and right homologous regions corresponding to part of the 3’-*UTR* were used to create donor vectors. These donors were transfected into the NF54::pCRISPR line to generate edited lines that were verified for appropriate integration at the target site by junctional PCR and sequencing of the amplified PCR product (Figure 7B). Western blot analysis of the AAT and HT lines showed aTc-dependent regulation of protein expression (Figure 7C). Despite in-frame integration of the epitope tag with FC, no tagged FC protein was detected under induced conditions. One possible explanation is that this protein is not highly expressed, which could be consistent with the low transcript levels observed across the intraerythrocytic developmental cycle (PlasmoDB.org: <u>PF3D7_1364900</u>).

**Figure 7.**
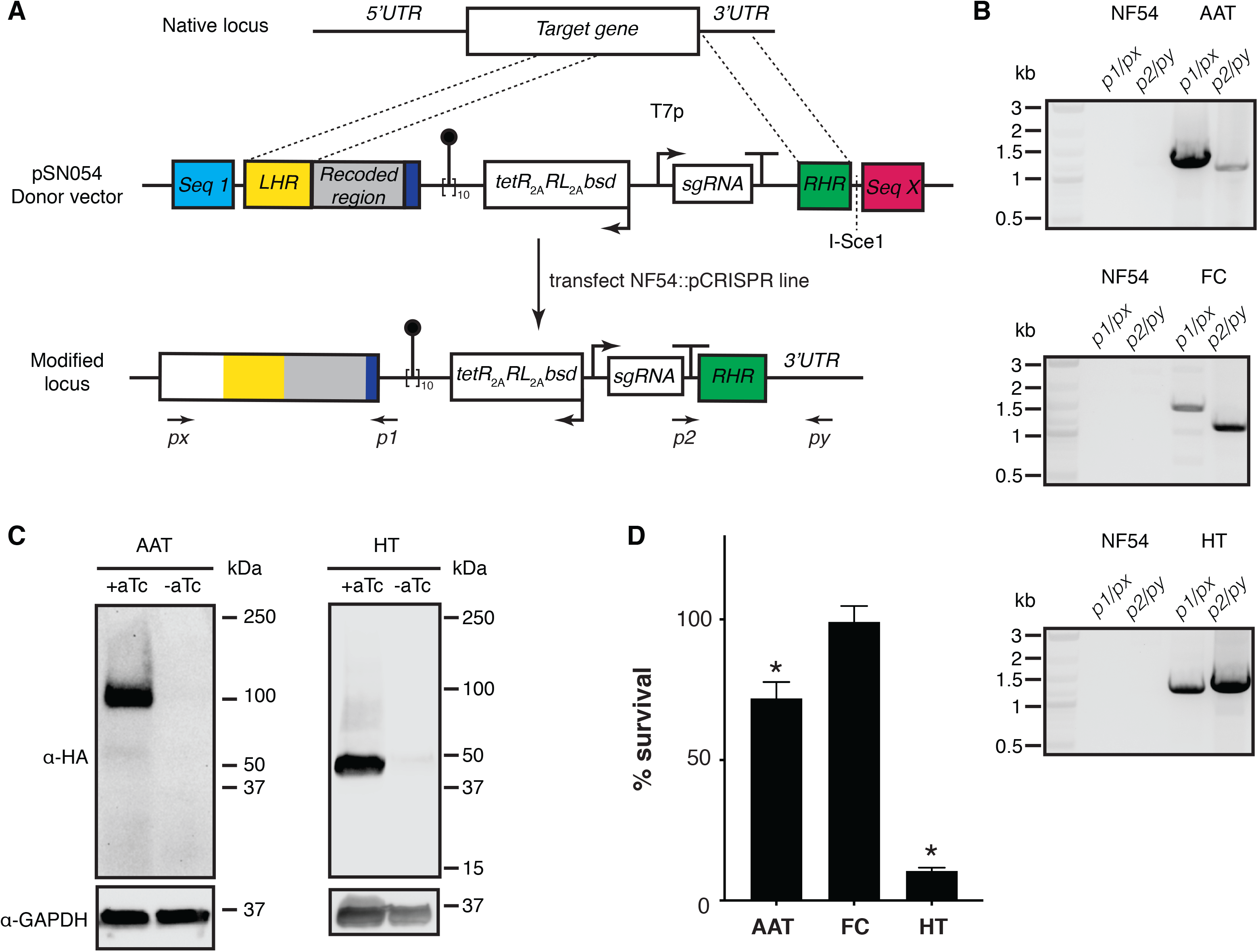
Manipulating the *3’-UTRs* of native loci using linear pSN054 donor vectors and CRISPR/Cas9 genome editing. (**A**) Generalized schematic of CRISPR/Cas9-mediated editing of a target locus to install C-terminal 2x-HA epitope tags, a 10x TetR aptamer array, and a TetR-DOZI, *Renilla* luciferase and Blasticidin S deaminase expression cassette. Donor vectors were transformed into the NF54::pCRISPR parasite line to modify targeted loci. (**B**) gDNA extracted from transgenic parasites analyzed by PCR demonstrated formation of the expected *5’*- and *3’*- integration junctions at each targeted locus. Expected junctional PCR product sizes (*5’*, *3’*): *AAT*, putative <u>a</u>mino <u>a</u>cid transporter (1.4 kb, 1.1 kb); FC, ferrochelatase (1.5 kb, 1.2 kb); and HT, hexose transporter (1.3 kb, 1.4 kb). Marker = 1 kb Plus DNA ladder (New England Biolabs). (**C**) Western blot analysis of target protein expression under ± aTc conditions. Expected molecular weights of the 2x HA-tagged proteins are *AAT* = 137.6 kDa and HT = 60.5 kDa. No HA-tagged FC protein was detected, even after multiple attempts. *Note*: AAT and HT migrate faster than expected based on their molecular weights; however, N- and C-terminal tagged HT migrate identically (Figure 5). (**D**) Normalized *Renilla* luciferase levels for parasites grown in 0 nM or 50 nM aTc for 72 h. Data represent the mean of *n = 3* ± s.e.m. * *p* ≤ 0.05 by Student’s t-test.

Inducing knockdown of HT and AAT, and putatively FC produced a range of responses in parasite growth. For HT, we detected a dramatic loss in parasite viability upon HT depletion, consistent with an essential role. This effect occurred in standard high glucose concentration (2 mg/mL) media, in contrast to the relatively more subtle glucose concentration-dependent growth phenotype observed with the regulated, synthetic promoter (Figure 7D and Supplementary Figure 3). This is consistent with the greater degree of protein knockdown achieved using a 10x aptamer array within the 3’-*UTR* (Figure 7C) compared to a single aptamer regulating the synthetic 5’-*UTR* (Figure 5D). For AAT, we observed an ∼30% decrease in parasite viability (Figure 7D) after a single replication cycle during which verifiable depletion of detectable protein occurred (Figure 7C). Interestingly, both *piggyBAC* mutagenesis (Zhang et al., 2018) and *P. berghei* knockout screens (Bushell et al., 2017) classify this gene as essential, although the knockdown studies here suggest that acute loss-of-function of this gene induces a fitness cost. Putative FC depletion through aTc removal did not result in any detectable change in parasite growth, which would be consistent with the non-essentiality of this protein and *de novo* heme biosynthesis during blood stage parasite development (Bushell et al., 2017; Ke et al., 2014; Nagaraj et al., 2013; Rathnapala et al., 2017; Zhang et al., 2018).

Even broader utility of pSN053/054 for enabling in-depth functional studies of diverse parasite genes is underscored by their application in elucidating roles for the claudin-like apicomplexan microneme protein (CLAMP) ligand in red blood cell invasion (Sidik et al., 2016), aspartate proteases Plasmepsin IX and X in red blood cell invasion and egress (Nasamu et al., 2017), and FtsH1 and ATG8 in apicoplast biogenesis/maintenance (Amberg-Johnson et al., 2017; Walczak et al., 2018). In some instances, it can be desirable to achieve even more stringently regulated knockdown to study biological function using a dual aptamer configuration (Ganesan et al., 2016). This is readily attained via pSN053/pSN054 (Supplementary Figure 1C), as illustrated during work defining new, essential roles for the integral membrane protein, EXP2 (Garten et al., 2018) and Plasmepsin V protease (Polino et al., 2018) beyond their previously described roles in protein export from the parasite to the red blood cell compartment (de Koning-Ward et al., 2016; 2009).

## CONCLUSIONS

We describe a flexible plasmid toolkit that improves the ease with which genome manipulation of the less genetically tractable but devastating human malarial parasite, *Plasmodium falciparum*, can be engineered to study gene function. This toolkit leverages the significantly increased efficiency linear plasmids afford in manipulating large and complex AT-rich DNA sequences without the undesirable deletions and rearrangements frequently encountered with typically-used circular bacterial plasmids. We have emphasized modular designs to allow facile and standardized vector configuration to address a broad variety of functional applications, including gene complementation, conditional regulation of gene expression, and gene deletions. We have ensured direct compatibility with CRISPR/Cas9 genome engineering by hardwiring an easily reprogrammed guide RNA production module into base donor vectors. We have also validated a pCRISPR plasmid that can be used to transiently or constitutively produce Cas9 when either co-transfected as a helper plasmid or stably integrated into a desired background strain, respectively. Importantly, any construct assembled in a linear vector can be ‘rescued’ with high efficiency to a circular BAC, while keeping AT-rich and repetitive sequences intact. In *P. falciparum*, only circular DNA is stably maintained episomally, and integrase-mediated site-specific integration is efficiently achieved from circular DNA. Therefore, this feature extends the approaches accessible for evaluating gene function in human malarial parasites. Altogether, we anticipate the toolkit described here will enable more technically robust strategies for high-resolution genome manipulation to improve our fundamental understanding of malaria parasite biology and enable applications that can enhance discovery of novel antimalarial therapeutics and disease prevention strategies.

## ACKNOWLEDGEMENTS

We thank Patrick Guye, Tharathorn Rimchala and Ron Weiss for helpful technical advice and sharing plasmid reagents. The NF54^attB^ cell line was a kind gift from Prof. David A. Fidock, Columbia University, New York. This research was supported by grants from the National Institutes of Health Directors New Innovator Award (1DP2OD007124, JCN); NIGMS Center for Integrative Synthetic Biology (P50 GM098792); NIGMS Biotechnology Training Grant (5-T32-GM08334, JCW); NIEHS Predoctoral Training Grant (5-T32-ES007020, BAW); and the Bill and Melinda Gates Foundation grants through the Grand Challenges Exploration initiative (OPP1132312, JCN), OPP1162467 (JCN) and OPP1158199 (JCN).

## AUTHOR CONTRIBUTIONS

All authors contributed to conceiving and designing experiments. ASN, AF, CFP, BAW and SMG performed experiments and analyzed data. JCW (pCRISPR plasmid and NF54::pCRISPR cell line), SJG (pSG372 plasmid) and SMG (pMG69) provided key reagents. ASN, AF, CFP, and JCN wrote the paper. JCN supervised the research.

## METHODS

### Molecular biology and plasmid construction

The methods used to assemble the vectors reported in this study are included as Supplementary Methods. These are described in step-by-step detail to allow users with basic knowledge of molecular cloning to easily build constructs in the pSN150, pSN154 and pSN053/054 contexts.

### Parasite culturing and transfection

*P. falciparum* strain 3D7 parasites were grown under 5% O_2_ and 5% CO_2_ in RPMI-1640 media supplemented with 5 g/L Albumax II (Life Technologies), 2 g/L sodium bicarbonate, 25 mM HEPES pH 7.4 (pH adjusted with potassium hydroxide), 1 mM hypoxanthine and 50 mg/L gentamicin. Transfections were performed using the red blood cell preloading method as described previously (Deitsch et al., 2001). Briefly, 50-100 μg of purified plasmid DNA were mixed with human red blood cells in 0.2 cm cuvettes and subjected to 8 square wave electroporation pulses of 365 V for 1 ms each, separated by 0.1 s. The DNA preloaded red blood cells were inoculated with schizont-stage parasites (e.g. NF54^attB^, NF54::pCRISPR) to achieve starting parasitemias ≤1% in RPMI 1640 Complete media. Beginning 4 days post-transfection, cultures were selected with either 2.5 µg/mL Blasticidin (RPI Corp, B12150-0.1) or a combination of 2.5 µg/mL Blasticidin and 2.5 nM WR99210 (Jacobus Pharmaceuticals). In knockout experiments performed on the NF54::pCRISPR background, parasites were selected with 1.5 µM DSM1 (MR4) only beginning on Day 4 post-transfection. For creating knockdown lines using pSN150 and pSN053/54 vectors, 500 nM anhydrotetracycline (aTc; Sigma-Aldrich, 37919) was included at the beginning of transfections and maintained throughout. Transfection progress was monitored using Giemsa-stained smears and *Renilla* luciferase measurements. High purity genomic DNA was isolated using the Blood & Cell Culture DNA Mini Kit (Qiagen 13323) according to the manufacturer’s instructions. Cultured cells were lysed under denaturing conditions, and Proteinase K was added to degrade proteins. The suspension was loaded onto a sterile Qiagen Genomic tip. After the recommended incubation period, the sample was centrifuged and the supernatant discarded. Following a wash step, the silica membrane was transferred to a new snap cap, safe lock microcentrifuge tube and distilled, deionized water or TE buffer used to elute DNA. Modification of the target locus was determined by PCR using appropriate locus-specific primers (Supplementary Methods).

### Western blotting analysis

Western blotting for *Sp*Cas9 and T7 RNAP was described previously (Wagner et al., 2014). To determine regulation of target protein expression by the TetR- or TetR-DOZI-aptamer system, parasites cultured with 0 nM or 50 nM aTc for 72 h were saponin-lysed, washed with 1x PBS, and proteins solubilized in lysis buffer containing 4% sodium dodecyl sulfate (SDS) and 0.5% Triton X-114 in 1x PBS. Protein extracts were mixed with loading buffer containing SDS and dithiothreitol (DTT) and loaded onto Mini-PROTEAN® TGX™ Precast Gels (4-15% gradient; Bio-Rad 4561084) in Tris-Glycine buffer. After separation by polyacrylamide gel electrophoresis (PAGE), proteins were transferred to a polyvinylidene fluoride (PVDF) membrane using the Mini Trans-Blot Electrophoretic Transfer Cell system (Bio-Rad) per the manufacturer’s instructions and blocked with 100 mg/mL skim milk in 1x TBS/Tween 20. PVDF membrane-bound proteins were first probed with mouse anti-HA (1:3000; Sigma, H3663) and rabbit anti-GAPDH (1:5000; Abcam, AB9485) primary antibodies and anti-mouse (1:5000; Thermo Fisher Scientific, 62-6520) and anti-rabbit (1:5000; Cell Signaling, 7074S) horseradish peroxidase (HRP)-conjugated secondary antibodies. Following incubation in SuperSignal® West Pico Chemiluminescent substrate (Thermo Fisher Scientific, PI34080), protein blots were imaged and analysed using the ChemiDoc™ MP System and Image Lab 5.2.0 (Bio-Rad).

### Luciferase and quantitative growth assays

Luciferase assays to track transfection progress and to measure aTc-dependent regulation of a firefly reporter were performed as previously described (Ganesan et al., 2016; Wagner et al., 2013) using the Dual-Luciferase Reporter® Assay System (Promega, E1910), Renilla Luciferase Assay System (Promega, E2810) or Renilla-Glo® Luciferase Assay System (Promega E2750) per manufacturer’s protocols. Quantitative growth assays were performed in 96-well U-bottom plates (Corning^®^ 62406-121) using synchronous ring-stage parasites set up in triplicate and cultured in the 0 nM or 50 nM aTc. Relative growth was determined using luciferase levels measured at 0 h (initial setup values) and after 72 h using the GloMax® Discover Multimode Microplate Reader (Promega). Luminescence values were normalized to samples treated with a lethal dose of chloroquine (200 nM) as no growth. Data was analyzed using GraphPad Prism (version 8; GraphPad Software). For GSK3 knockdown experiments, parasite growth was determined by FACS analysis using an Accuri Flow Cytometer (BD Biosciences). Parasites were stained for nucleic acid content with 1 μM SYTO 61 (Life Technologies) and analyzed in the FL4 signal channel to determine the fraction of parasitized red blood cells (parasitemia).

For comparing the functional HT regulation achieved using the pSN150 and pSN054 configurations, we measured the expansion of synchronous ring stage parasites after a 72 h period of growth. Assays were set up at various initial parasitemias (0.1%, 0.5% and 1.5%) in various glucose (0.2, 0.8, 1.4 and 2.0 mg/mL) and aTc (0, 1, 3 and 50 nM) concentrations. RPMI 1640 without glucose (US Biological, R8999-13) was used as the base medium, with supplementation as needed with glucose and aTc.

## SUPPLEMENTARY FIGURES

**Supplementary Figure 1.**
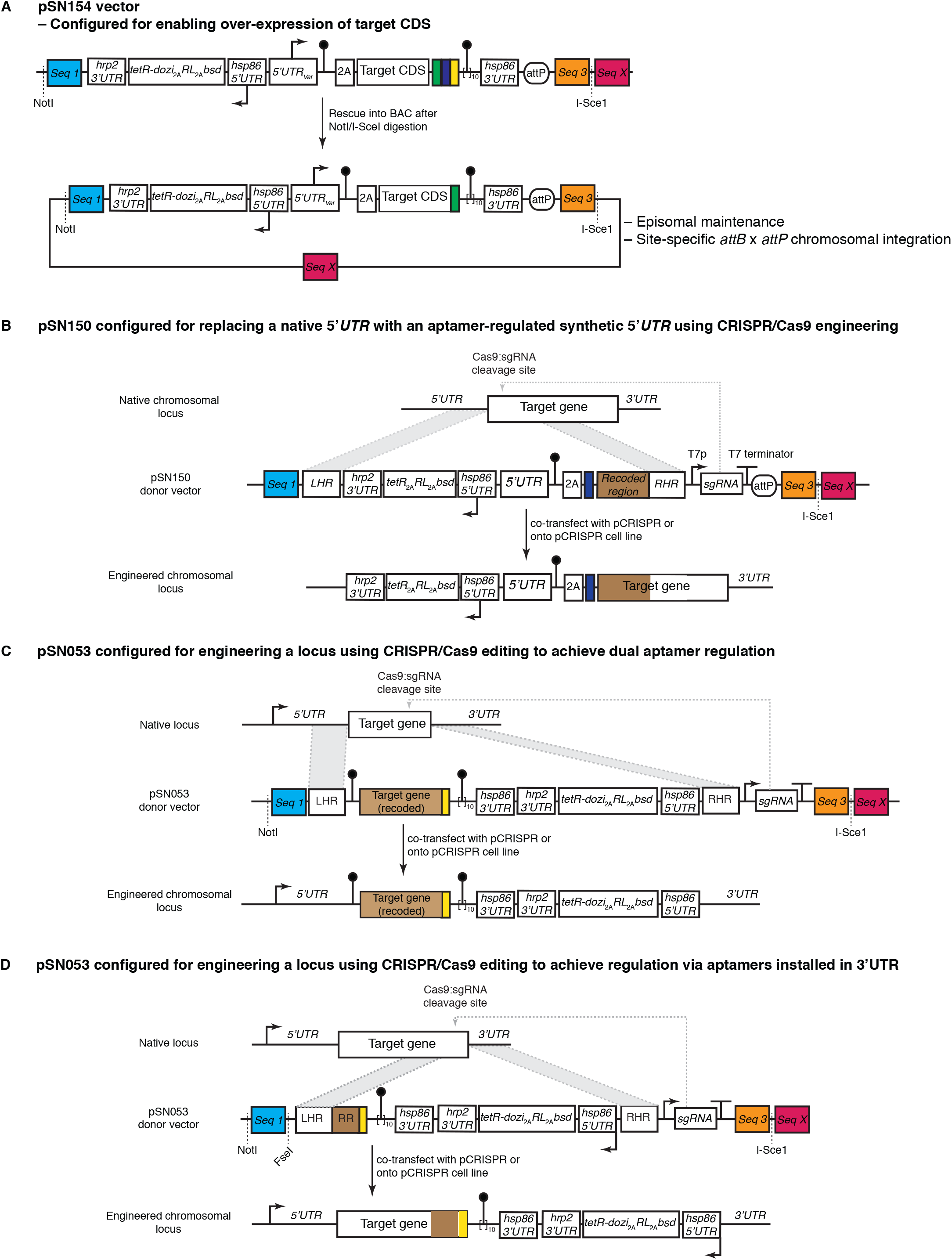
Several operations using the pSN vector family for achieving TetR aptamer-regulated conditional regulation of gene expression after genome editing are summarized. (**A**) Expressing a transgene from pSN154 rescued into a BAC that is then either stably maintained as an episome or integrated at a chromosomal *attB* site using the *Bxb1* integrase. (**B**) Using pSN150 and CRISPR-Cas9 genome editing to swap a native promoter for a synthetic promoter regulated by a single TetR aptamer in the 5’-*UTR* to conditionally regulate expression of an endogenous gene from its native chromosomal locus. (**C**) Using pSN053/54 and CRISPR/Cas9 genome editing to place a gene transcribed by its native promoter under dual TetR aptamer (5’- and 3’-*UTR* located) conditional regulation by TetR-DOZI. (**D**) Using pSN053/54 and CRISPR/Cas9 genome editing to place a gene transcribed by its native promoter under conditional regulation by TetR-DOZI via 3’-*UTR* located TetR aptamers.

**Supplementary Figure 2.**
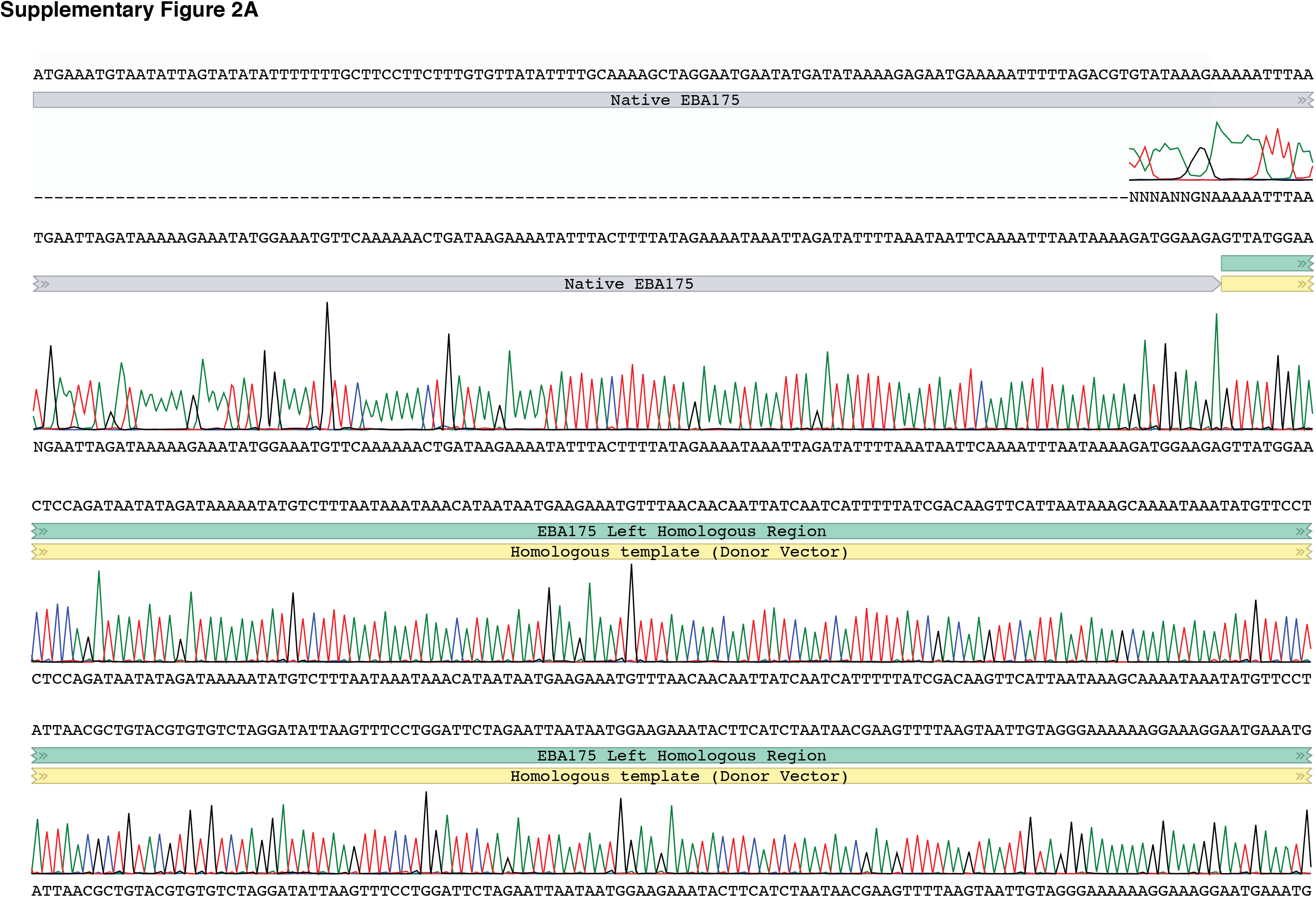
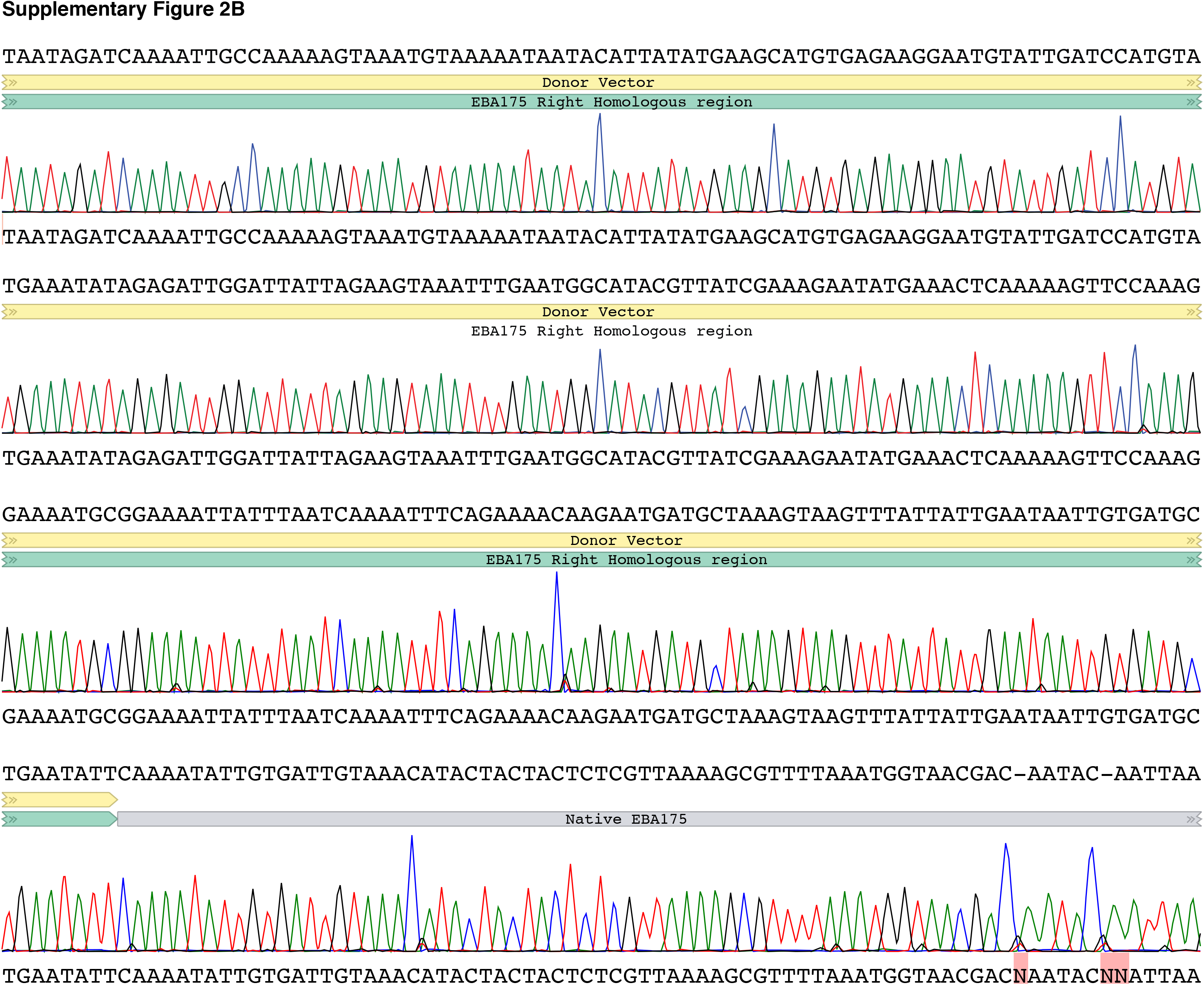
Complete Sanger sequencing traces of the (**A**) 5’- and (**B**) 3’-junction integration PCR products from the *eba175* knockout parasite line.

**Supplementary Figure 3.**
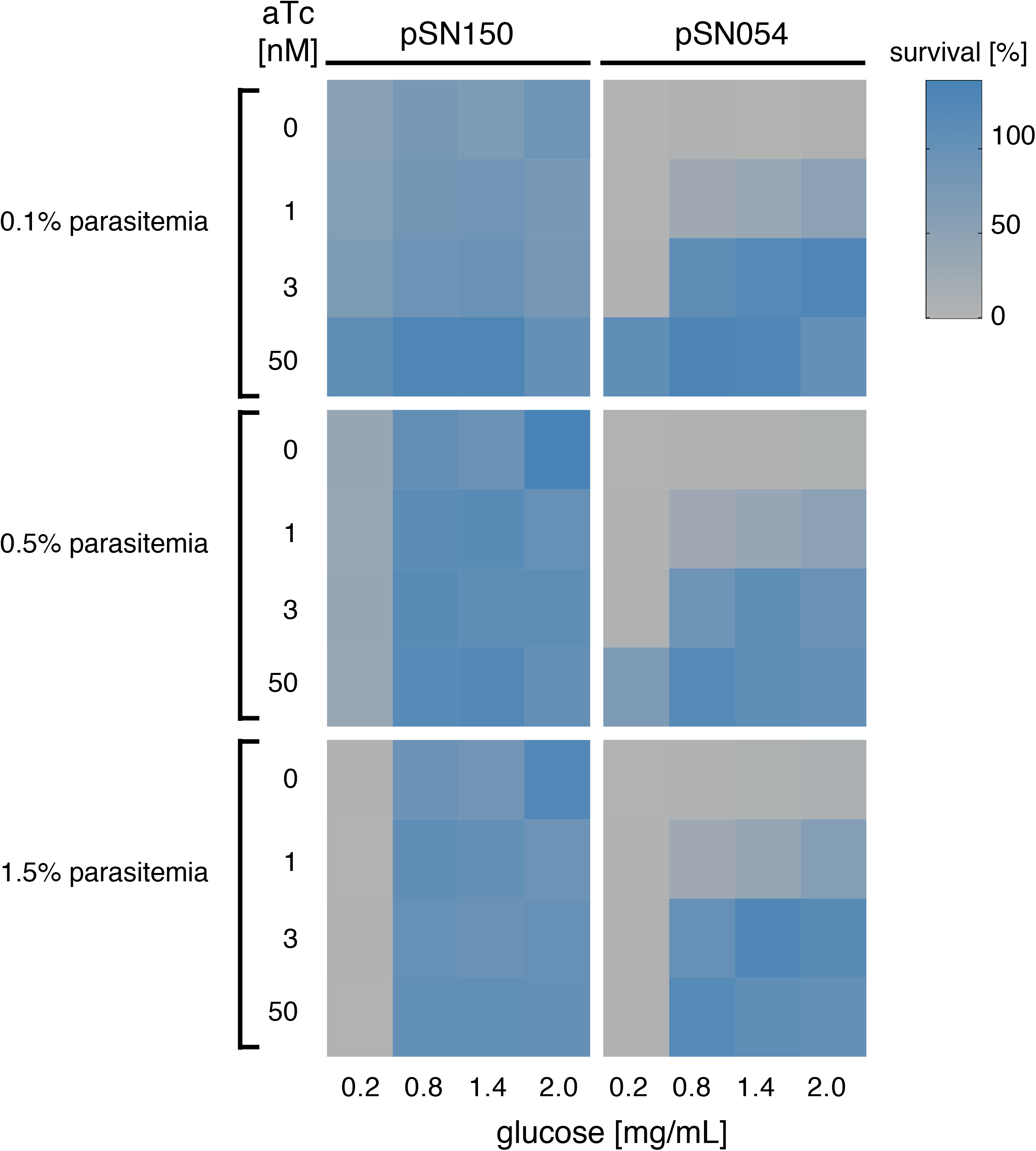
Growth of parasites in which the HT locus has been engineered to achieve regulated expression via a non-native, synthetic promoter/*5’-UTR* regulated by a single TetR aptamer within the transcript’s *5’-UTR* (pSN150) or its native promoter and a 10x aptamer array positioned within the *3’-UTR* of the transcript (pSN054). The relative growth of ring stage parasites inoculated at varying initial parasitemia (0.1%, 0.5% and 1.5%) in media containing a range of glucose (0.2, 0.8, 1.4 and 2.0 mg/mL) and aTc (0, 1, 3 and 50 nM) concentrations was determined after 72 h. Parasite growth was determined by monitoring *Renilla* luciferase levels, with the values obtained at 2 mg/mL glucose and 50 nM aTc condition representing maximal (100%) growth. Data are the mean ± s.d. from one of two independent experiments.

## SUPPLEMENTARY METHODS

### Materials

pJAZZ-OC Not I Vector (Lucigen, Catalog # 43024)

BigEasy-TSA Electrocompetent Cells (Lucigen, Catalog # 60224)

BAC-Optimized Replicator™ v2.0 Electrocompetent cells (Lucigen, Catalog # 60210)

2x Gibson Assembly Master Mix (NEB, Catalog #E5510)

DNA Polymerase I, Large (Klenow) Fragment (New England Biolabs, Catalog# M0210)

Restriction enzymes were purchased from New England Biolabs, unless otherwise stated

### Section I. Creating base linear vectors while incorporating components from traditional circular plasmids

The pJAZZ-OC vector (Lucigen) was converted into the various pSN configurations as summarized in the scheme below (*Supplementary Methods Figure 1*).

**Supplementary Methods Figure 1.**
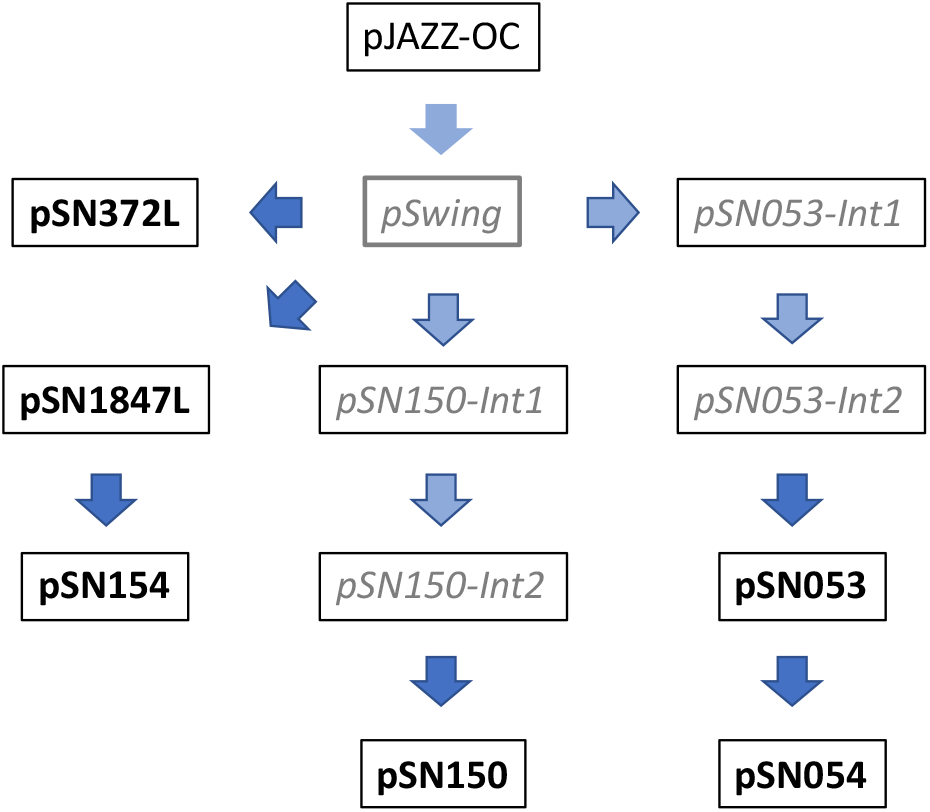
A flow chart summarizing conversion of pJAZZ-OC into the various linear pSN base vectors. Grey italics and light blue arrows denote intermediate steps en route to final plasmids (boldface and darker blue arrows).

### pSwing construction

pJAZZ-OC Not I Vector (200 ng) was digested with NotI at 37 °C for 90 min and the reaction heat inactivated at 65 °C for 20 min. The pSwing gblock (*Supplementary Methods Figure 2*) containing the unique sequences 1, 3 and X (SEQ1, SEQ3, SEQX) and FseI, SacII and I-SceI restriction sites was obtained from IDT. The digested vector (20 ng) and gblock (20 ng) were mixed with an equal volume of 2x Gibson Assembly Master Mix and incubated at 50 °C for 1 h to assemble an intermediate vector, pSwing. Big Easy TSA cells were transformed with 1 µL Gibson reaction mixture, and plated on LB-agar with chloramphenicol (34 µg/mL) and incubated overnight at 30 °C. Selected colonies were grown overnight in liquid LB supplemented with chloramphenicol (34 µg/mL), mini-prepped and verified by restriction digestion and sequencing.

**Supplementary Methods Figure 2.**
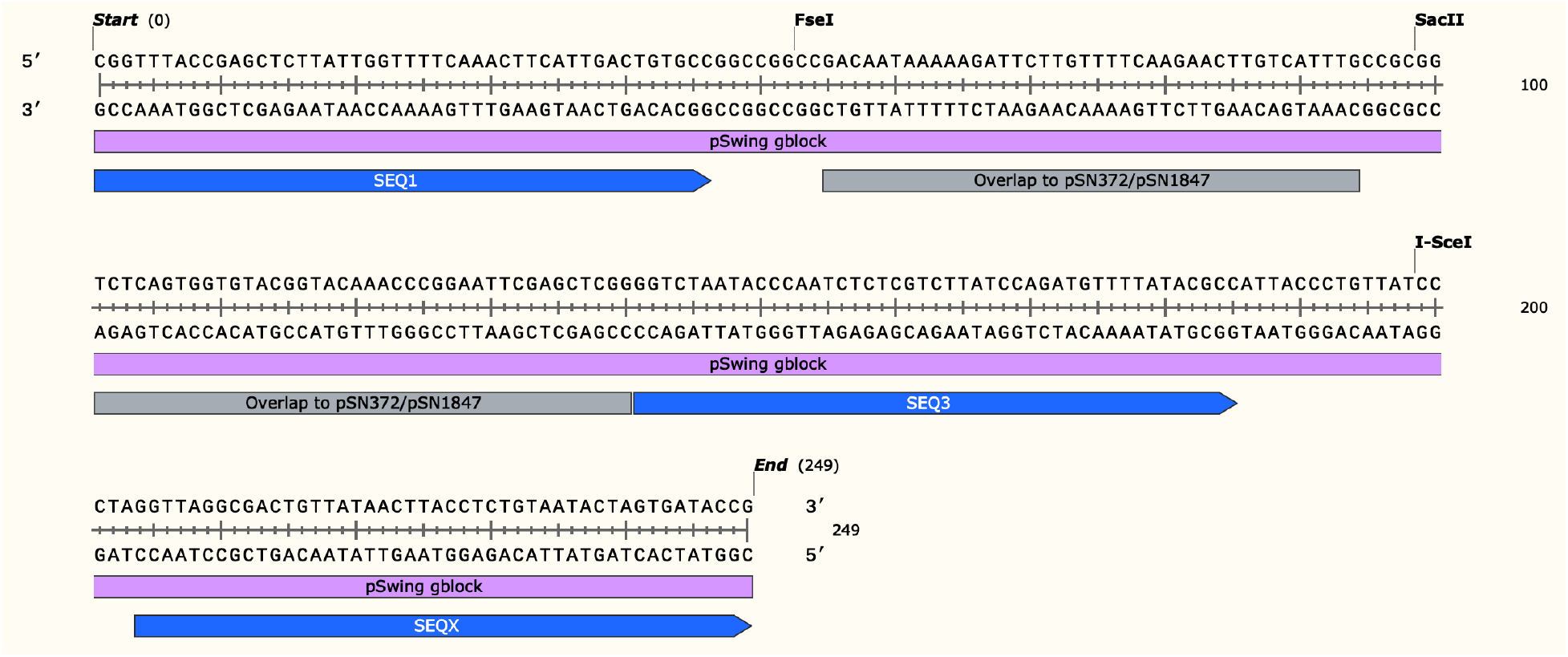
pSwing gblock sequence map and features.

### pSN372L and pSN1847L construction

Plasmids pSG372.3 and pMG56 (Ganesan *et al*, 2016) were first modified to produce pSN372 and pSN1847. pSN372 was made by inserting via Gibson assembly: (1) a loxP-containing DNA fragment (<u>tctattattaaataaatttaatgga</u>**ataacttcgtatagcatacattatacgaagttat**<u>ccggtttagccctcccacacataac</u>; loxP site in bold font and Gibson overlap underlined) via an AgeI site upstream of BSD; and (2) a second loxP-containing fragment (<u>atacctaatagaaatatatcttaag</u>**ataacttcgtatagcatacattatacgaagttat***aataaatacctaatacaatccaggc*<u>caatcca</u> <u>ggcagagaaaggtcgatac</u>; loxP site in bold, spacer in italics and Gibson overlap underlined) via an AflII site downstream of the Cam promoter. pSN1847 was created by performing the second step above on pMG56. Linear pSN372L and pSN1847L were created from their respective pSN372 and pSN1847 circular plasmids. pSwing (200 ng) was digested with SacII at 37 °C for 1.5 h, followed by heat inactivation of enzyme at 65 °C for 20 min. Inserts from pSN372 and pSN1847 were released by ScaI+NotI double digestion at 37 °C for 1 h, followed by heat inactivation of enzymes at 80 °C for 20 min. Released inserts (20 ng) were each mixed with SacII-digested pSwing (20 ng) and an equal volume 2x Gibson Assembly Master Mix, and incubated at 50 °C for 1 h. Big Easy TSA cells were transformed and colonies analyzed as above to isolate pSN372L and pSN1847L (*Supplementary Methods Figure 3A,B*). Mini-prepped plasmids were analyzed for proper assembly by NotI and FseI+AflII (pSN372L) and Aflll+NotI (pSN1847L) digestions.

**Supplementary Methods Figure 3.**
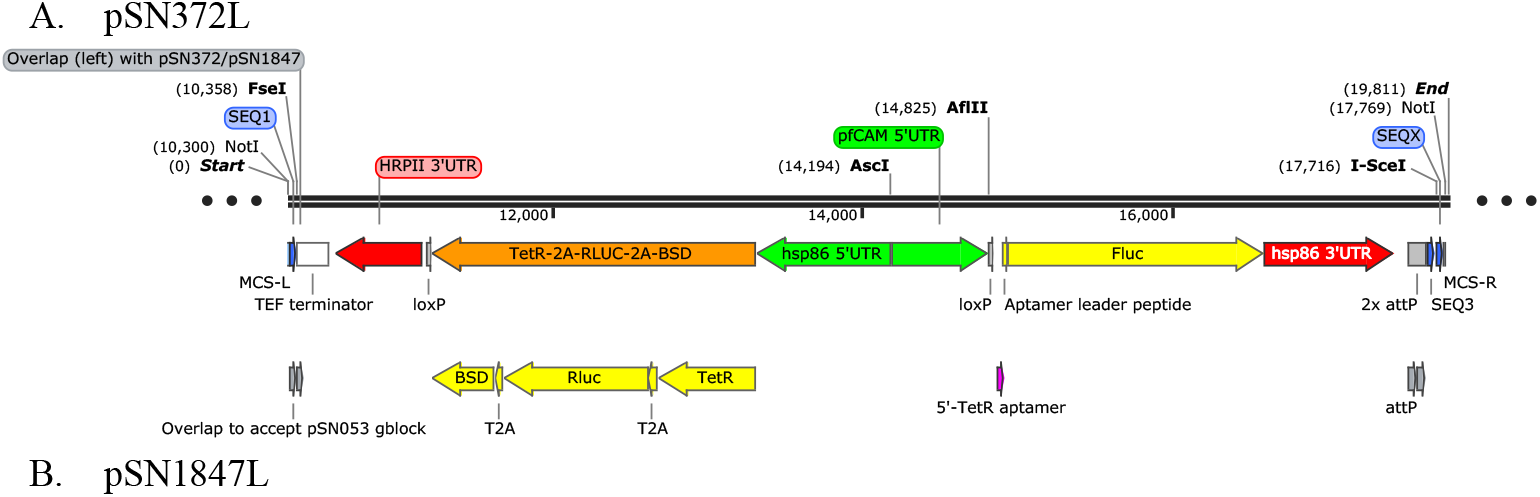
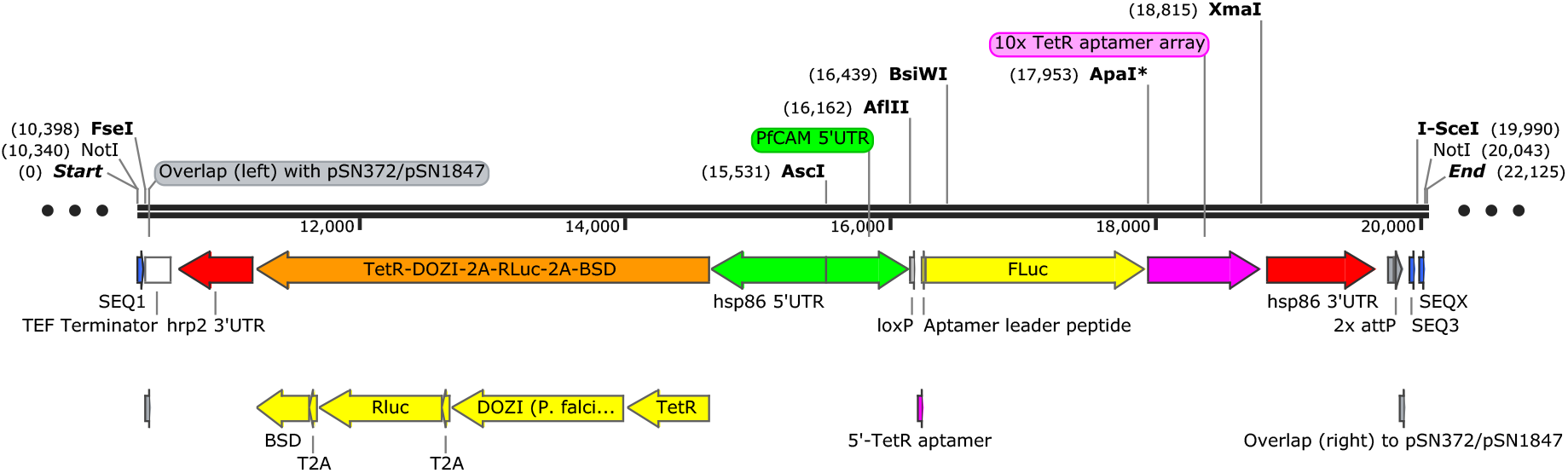
Overview map of (A) pSN372L and (B) pSN1847L.

### pSN154 construction

The *Firefly* luciferase gene in pSN1847L was removed by AflII+ApaI digestion and replaced with the pSN154-gblock to obtain pSN154 (*Supplementary Methods Figure 4A,B*).

**Supplementary Methods Figure 4.**
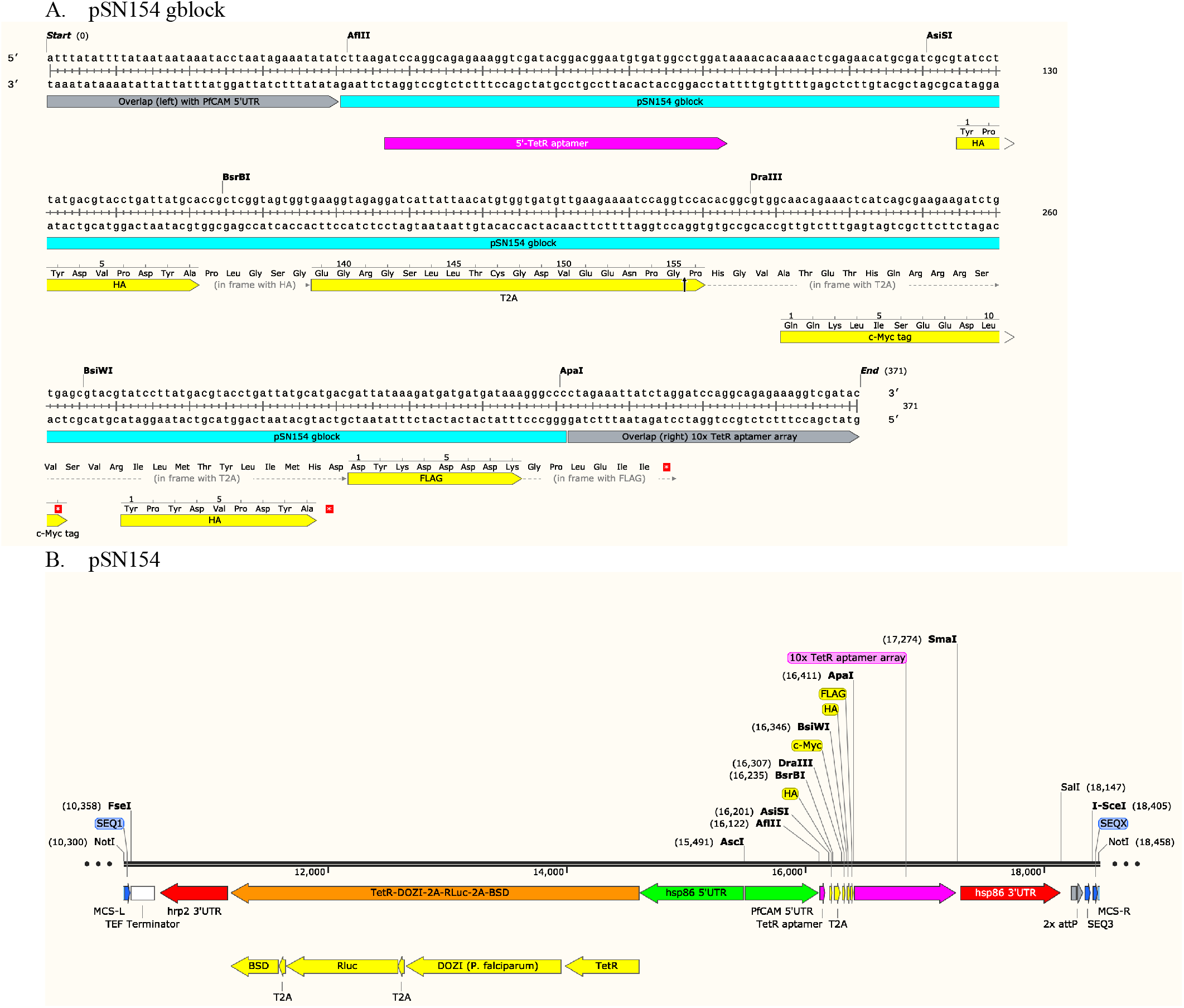
Overview of pSN154 construction. (A) pSN154-gblock sequence and features. (B) Map of final pSN154 plasmid.

### pSN150 construction

The pSN150-gblock (IDT) was cloned by Gibson assembly into FseI+SacII-digested pSwing to create pSN150-Int1 (*Supplementary Methods Figure 5A,B*). A SEQ1-(FseI)-*hrp2* 3’UTR-TetR-2A-RLuc-2A-BSD-*hsp86* 5’UTR-(AscI)-*PfCAM* 5’UTR-loxP-TetR aptamer fragment released by NotI+XhoI digestion of pSN372L was cloned by Gibson assembly into FseI-digested pSN150-Int1 to produce pSN150-Int2 (*Supplementary Methods Figure 5C*). Lastly, an sgRNA gblock (IDT) consisting a [5’-cMyc tag]-[T7 promoter]-(I-ppoI/AflII restriction site)-[sgRNA scaffold]-[T7 terminator]-[3’-HA tag] was installed by Gibson assembly into BsiWI-digested pSN150-Int2 to create pSN150 (*Supplementary Methods Figure 5D*).

### pSN053 and pSN054 construction

The pSN053-gblock was cloned by Gibson assembly into FseI+SacII-digested pSwing to yield pSN053-Int1 (*Supplementary Methods Figure 6A,B*). A fragment containing the *hsp86* 5’*UTR*-TetR-DOZI_2A_-RLuc_2A_BSD-*hrp2* 3’*UTR* transcription unit was released from pSN1847 by NotI+SacI digestion, and inserted by Gibson assembly into ApaI-digested pSN053-Int1 to yield pSN053-Int2 (*Supplementary Methods Figure 6C*). Lastly, a fragment containing c-Myc-HA-FLAG-10x TetR aptamer-*hsp86* 3’UTR was released from pSN154 by DraIII+SalI digestion and installed by Gibson assembly into SacII-digested pSN053-Int2 to yield pSN053 (*Supplementary Methods Figure 6D*).

To obtain pSN054, I-SceI-digested pSN053 was Gibson assembled with pSN054-conversion gblock to yield pSN054. Thus, pSN054 differs from pSN053 in that the I-SceI site is immediately downstream of SEQ3 and SEQX is deleted (*Supplementary Methods Figure 6E*).

**Supplementary Methods Figure 5.**
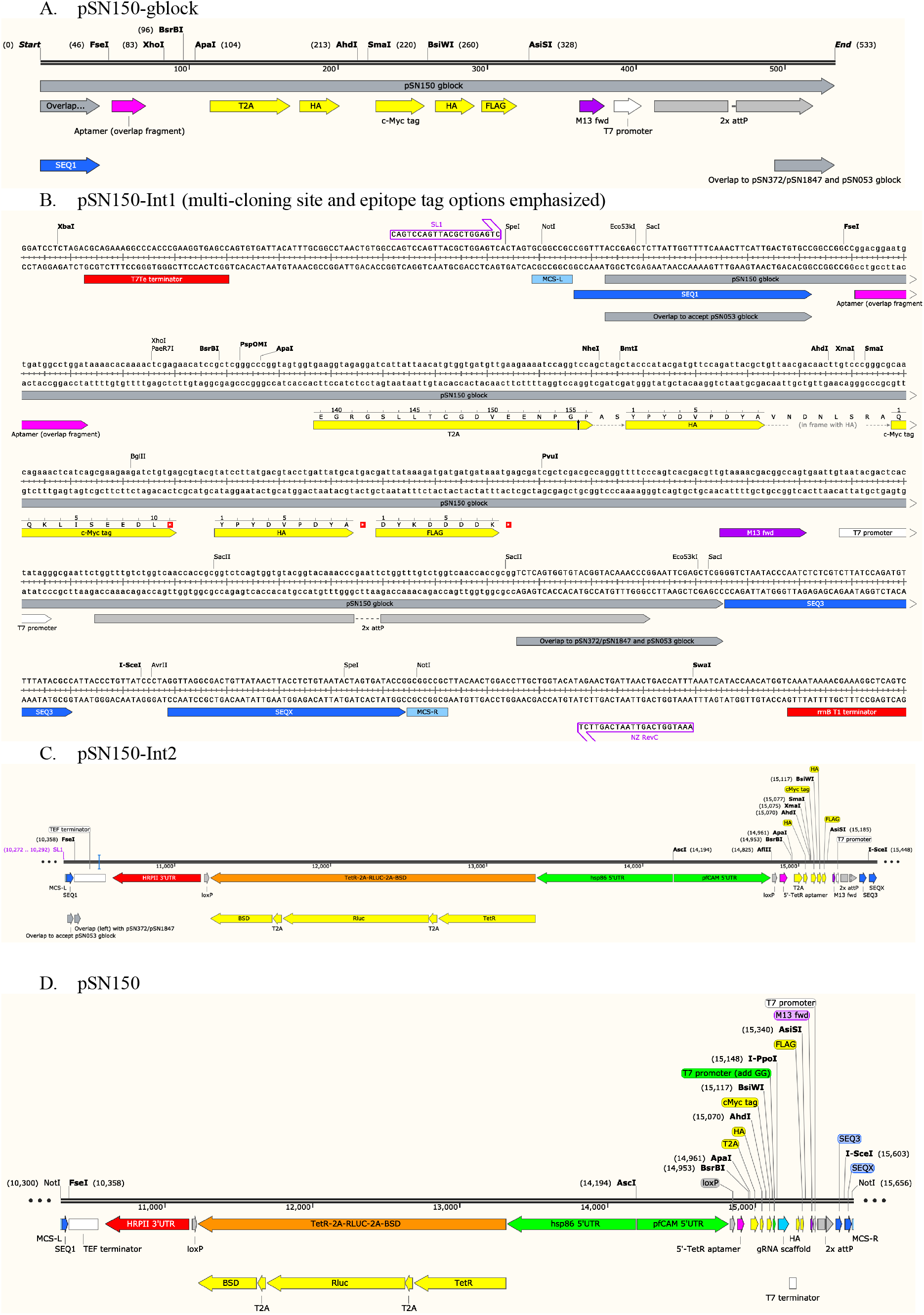
Overview of pSN150 construction. Key features are shown for (A) pSN150 gblock, (B) pSN150-Int1, (C) pSN150-Int2 and (D) pSN150.

**Supplementary Methods Figure 6.**
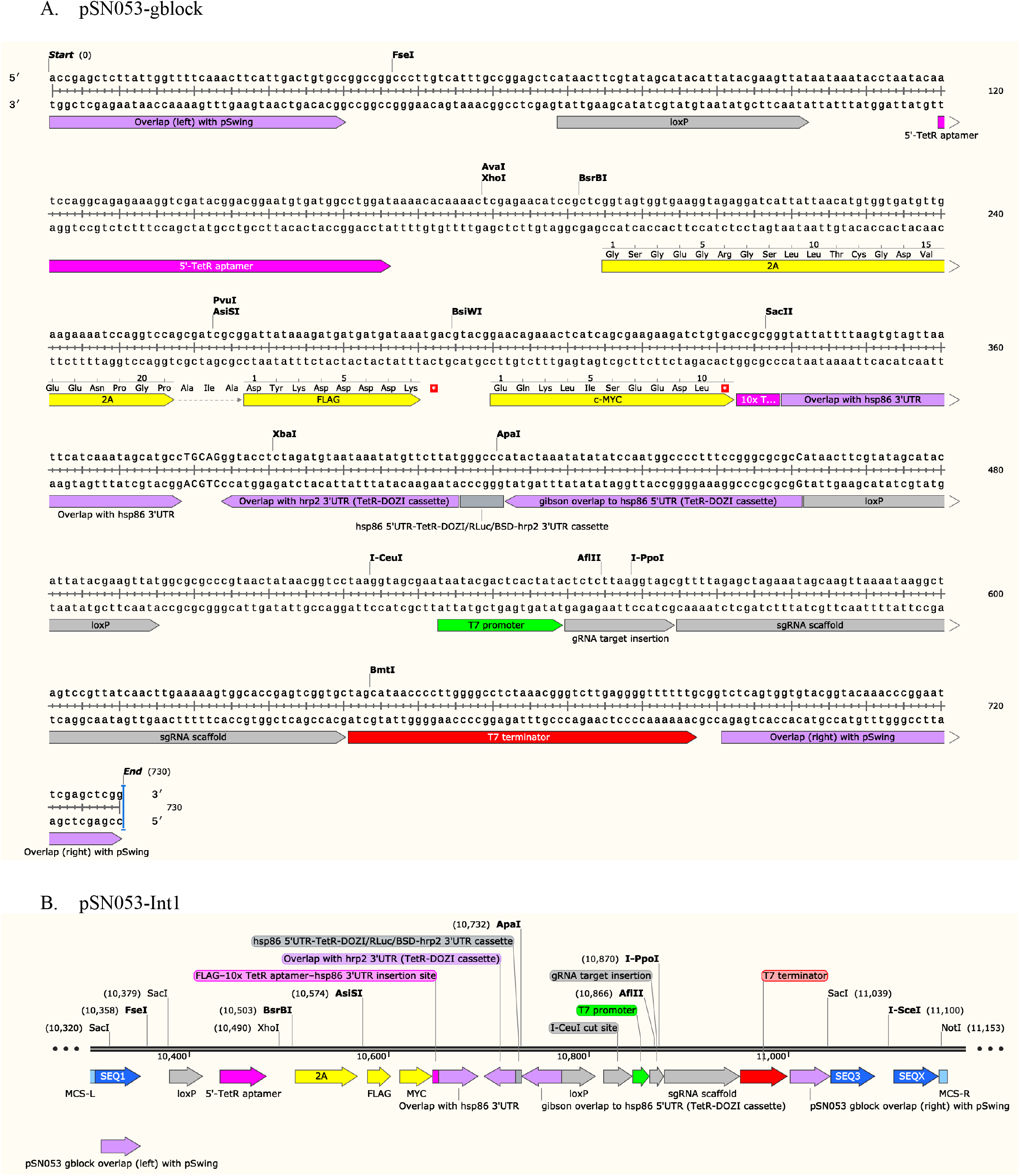
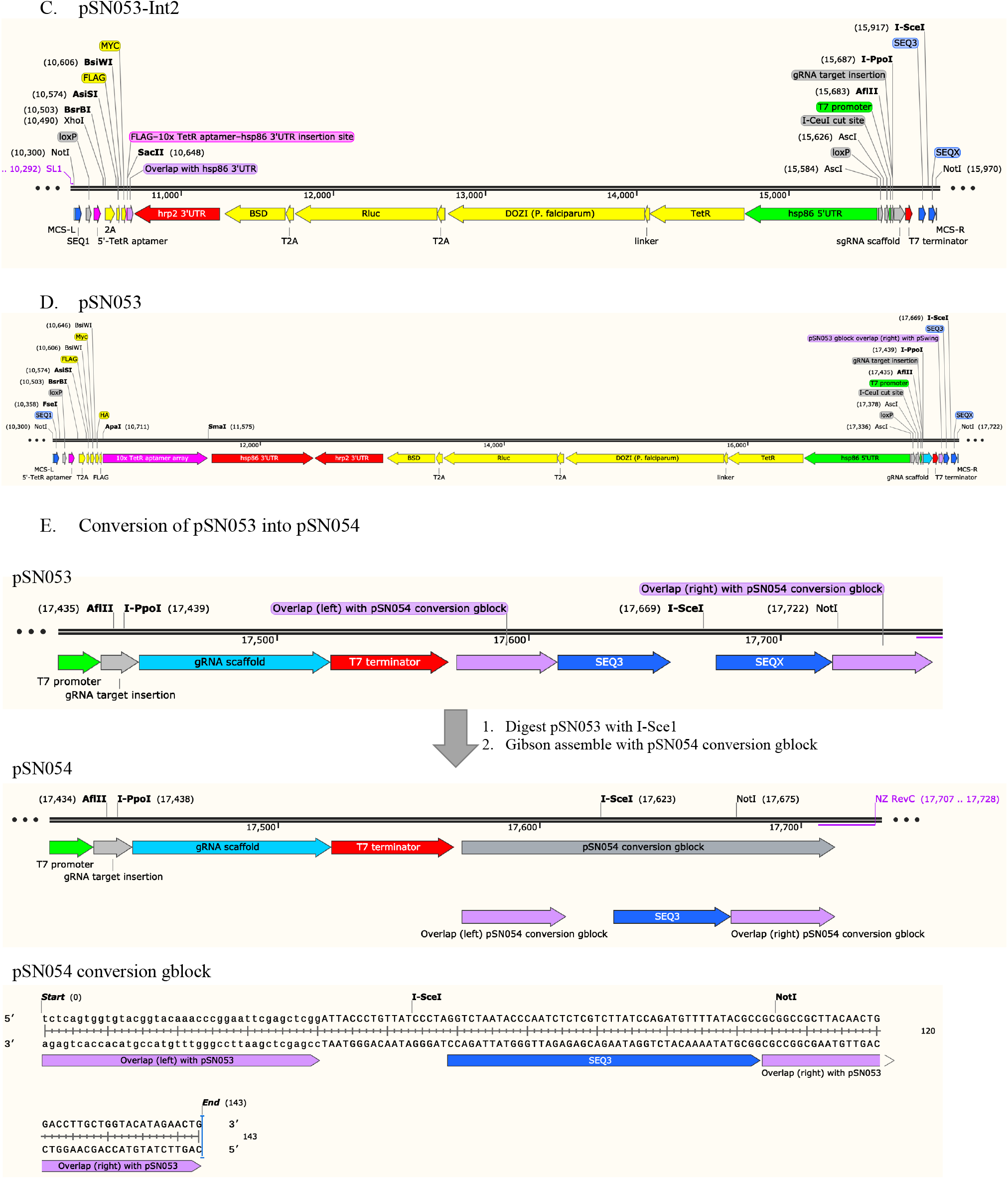
Overview of pSN053/4 construction. Key features are shown for (A) pSN053 gblock, (B) pSN053-Int-1, (C) pSN053-Int2, (D) pSN053 and (E) pSN053 conversion to pSN054.

### Section 2. Detailed methods for configuring each linear base vector (pSN154, pSN150, pSN053/pSN054) to achieve diverse locus modification outcomes in *P. falciparum*

**A. Assembling complementation/over-expression vectors for episomal maintenance in pSN154** *(Estimated time: 7 days)*

Day 1

1. Digest 200 ng pSN154 plasmid with the appropriate enzyme depending on one’s choice of epitope tag/tags (see table 1) in 10 µL at 37 °C for 90 min, followed by heat inactivation of enzymes at the appropriate temperature.

| Restriction enzymes | Tag on N-terminus | Tag on C-terminus |
| --- | --- | --- |
| AsiSI only | - | HA+T2A |
| AsiSI+BsrBI | - | T2A |
| AsiSI+DraIII; BsrBI+DraIII;<br>DraIII | - | cMyc, HA or FLAG |
| AsiSI+BsiWI; BsrBI+BsiWI;<br>DraIII+BsiWI | - | HA or FLAG |
| BsrBI | HA | T2A sequence or none if desired |

2. PCR the full-length gene of interest from genomic DNA (gDNA) or complementary DNA (cDNA) of the *P. falciparum* strain being studied with a high-fidelity polymerase. Gel purify the PCR product. Resuspend to 20 ng/µL in water.

3. Mix 20 ng each of PCR product and digested vector with 3 µL 2x Gibson Master Mix and incubate reaction at 50 °C for 1 h.

4. Transform Big Easy TSA cells with 1 µL Gibson reaction mixture Transformation conditions: 1 mm cuvette, 10 µF, 600 Ohms, 1800 V.

5. Plate on LB-agar with chloramphenicol (34 µg/mL) and incubate overnight at 30 °C. [**Note**: Big Easy TSA cells are ampicillin resistant.]

Day 2

Pick 8 colonies and grow overnight at 30 °C in liquid LB media containing chloramphenicol (34 µg/mL) and 0.2% w/v arabinose.

Day 3

1. Evaluate insertion of the gene of interest with an AflII and ApaI double digest.
2. Confirm correctly digesting plasmids by sequencing the inserted gene and its flanking sequences.

Day 4

1. Digest 200 ng of the linear plasmid with NotI and I-SceI in a 10 µL volume at 37 °C for 90 min followed by heat inactivation of enzymes at 80 °C.
2. Digest 200 ng of pBigBOB with PacI and heat inactivate enzyme at 65 °C for 20 min.
3. Digest 200 ng of pAdapter with XhoI and XbaI at 37 °C for 3 h and gel purify the plasmid backbone. Resuspend to 20 ng/µL. **NOTE:** pAdapter confers kanamycin resistance and XhoI cannot be heat inactivated.
4. Incubate 20 ng of each of the digested plasmid containing gene of interest, pBigBOB and pAdapter in a Gibson reaction at 50 °C for 1 h.
5. Transform BAC-Optimized Replicator™ v2.0 Electrocompetent cells with 1 µL Gibson reaction mixture. Transformation conditions: 1 mm cuvette, 10 µF, 600 Ohms and 1800 V.
6. Plate on LB-agar with chloramphenicol (34 µg/mL) and kanamycin (50 µg/mL) and incubate overnight at 30 °C.

Day 5

Pick 8 colonies and grow overnight at 30 °C in liquid LB media containing chloramphenicol (34 µg/mL) and kanamycin (50 µg/mL).

Day 6

1. Pick colonies and mini-prep plasmid DNA.
2. Digest with AvrII, PvuI and NotI+XhoI to confirm that the relevant cassettes are present.

**B. Assembling vectors to engineer promoter and *5’-UTR* regions of *P. falciparum* loci using pSN150** *(Estimated time: 10 days)*

Day 1: Installing recoded coding sequence (CDS) and right homologous regions (RHR).

1. Digest 200 ng pSN150 plasmid with the desired enzymes below in 10 µL at 37 °C for 90 min, followed by heat inactivation of restriction enzyme(s) at 80 °C for 20 min.

| Restriction enzymes | Tag on N-terminus |
| --- | --- |
| BsrBI+AhdI/BsiWI | None |
| AhdI; AhdI+BsiWI | HA |
| BsiWI | HA or cMyc |

2. PCR the RHR from gDNA using a high-fidelity polymerase and gel purify the product. Resuspend to 20 ng/µL.

3. Resuspend synthetic DNA corresponding to recoded CDS region in water to 20 ng/µL.

4. Mix 20 ng each of the RHR, gblock and digested vector with an equal volume 2x Gibson Assembly master mix. Incubate at 50 °C for 1 h.

5. Transform BigEasy-TSA Electrocompetent Cells with 1 µL Gibson reaction mixture. Transformation conditions: 1 mm cuvette, 10 µF, 600 Ohms, 1800 V.

6. Plate on LB-agar with chloramphenicol (34 µg/mL) and incubate overnight at 30 °C.

Day 2

Pick 8 colonies and grow overnight at 30 °C in liquid LB media containing chloramphenicol (34 µg/mL) and 0.2% w/v arabinose.

Day 3

1. Mini-prep plasmid DNA and test for insertion of both RHR and recoded DNA segment by restriction digestion with AflII and BsiWI.
2. Confirm correctly digesting plasmids by DNA sequencing and select the pSN150-RHR intermediate for the next step.

Day 4: Installation of the left homologous region (LHR)

1. Digest 200 ng pSN150-RHR plasmid with FseI in 10 µL at 37 °C for 90 min followed by heat inactivation of enzyme at 80 °C for 20 min.
2. PCR the LHR from gDNA using a high-fidelity polymerase and gel purify the product. Resuspend to 20 ng/µL.
3. Mix 20 ng LHR and digested vector with 3 µL 2x Gibson Assembly master mix, and incubate at 50 °C for 1 h.
4. Transform Big Easy TSA cells with 1 µL Gibson reaction mixture.
5. Plate on LB-agar with chloramphenicol (34 µg/mL) and incubate overnight at 30 °C.

Day 5

Pick 8 colonies and grow overnight at 30 °C in liquid LB media containing chloramphenicol (34 µg/mL) and 0.2% w/v arabinose.

Day 6

1. Mini-prep plasmid DNA and analyze colonies for LHR insertion NotI+AscI digest and RHR/recoded region retention AflII+BsiWI digest.
2. Confirm correctly digesting plasmids by sequencing the LHR and select pSN150-LHR-RHR intermediate for the next step.

Day 7: Installation of sgRNA into pSN150-LHR-RHR intermediate.

1. Digest 200 ng of the vector with I-ppoI or AflII.
2. Obtain sgRNA-encoding DNA fragment using complementary primers in a Klenow reaction (see protocol below) or purchase as synthetic DNA fragment. (*Optional*: The Klenow fragment can be agarose gel purified, if needed).
3. Mix 20 ng each sgRNA-encoding DNA and digested pSN150-LHR-RHR vector with 3 µL 2x Gibson Assembly master mix and incubate at 50 °C for 1 h.
4. Transform BigEasy-TSA Electrocompetent Cells with 1 µL Gibson reaction mixture. Transformation conditions: 1 mm cuvette, 10 µF, 600 Ohms, 1800 V.
5. Plate on LB-agar with chloramphenicol (34 µg/mL) and incubate overnight at 30 °C.

Day 8

Pick 8 colonies and grow overnight at 30 °C in liquid LB media containing chloramphenicol (34 µg/mL) and 0.2% w/v arabinose.

Day 9

1. Mini-prep plasmid DNA and analyze colonies for sgRNA-encoding DNA insertion by AflII (or I-ppoI) digest.
2. Sequence sgRNA insert using the T7 promoter or other suitable primer.
3. Confirm retention of inserted LHR and RHR regions using (NotI+AscI) and AflII+BsiWI restriction digests, respectively, in selected final colonies.

Day 10

Maxi-prep plasmid for *P. falciparum* transfection. Plasmids are quite stable during this step. However, as prudent practice, we recommend performing restriction digests as in the previous step to verify the absence of gross deletions/rearrangements.

**C. Assembling donor vectors to achieve gene deletions in *P. falciparum* using pSN150** *(Estimated time: 10 days)*

Day 1: Installation of left homologous region (LHR)

1. Digest 200 ng pSN150 plasmid with FseI in 10 µL volume at 37 °C for 90 min, followed by heat inactivation of enzyme at 80 °C for 20 min.
2. PCR the LHR from gDNA using a high-fidelity polymerase and gel purify the product. Resuspend to 20 ng/µL.
3. Mix 20 ng each of LHR and digested vector with 3 µL 2x Gibson Assembly master mix, and incubate at 50 °C for 1 h.
4. Transform BigEasy-TSA Electrocompetent Cells with 1 µL Gibson reaction mixture. Transformation conditions: 1 mm cuvette, 10 µF, 600 Ohms, 1800 V.
5. Plate on LB-agar with chloramphenicol (34 µg/mL) and incubate overnight at 30 °C.

Day 2

Pick 8 colonies and grow overnight at 30 °C in liquid LB media containing chloramphenicol (34 µg/mL) and 0.2% w/v arabinose.

Day 3

1. Mini-prep plasmid DNA and test for LHR insertion by restriction digestion with FseI+XbaI (or XbaI alone). [**Note**. Even though the XbaI site is not unique in this plasmid, digesting with this enzyme will produce a pattern unambiguously identifying LHR insertion.]
2. Confirm correctly digesting plasmids by sequencing the LHR, and select the pSN150-LHR intermediate for the next step.

Day 4: Installation of right homologous region (RHR)

1. Digest 200 ng pSN150-LHR plasmid with AscI+XmaI in a 10 µL at 37 °C for 90 min, followed by heat inactivation of enzymes at 80 °C for 20 min.
2. PCR the RHR from gDNA using a high-fidelity polymerase and gel purify the product. Resuspend to 20 ng/µL.
3. Mix 20 ng each of the RHR fragment and digested vector pSN150-LHR with 3 µL 2x Gibson Assembly master mix at 50 °C for 1 h.
4. Transform BigEasy-TSA Electrocompetent Cells with 1 µL Gibson reaction mixture. Transformation conditions: 1 mm cuvette, 10 µF, 600 Ohms, 1800 V.
5. Plate on LB-agar with chloramphenicol (34 µg/mL) and incubate overnight at 30 °C.

Day 5

Pick 8 colonies and grow overnight at 30 °C in liquid LB media containing chloramphenicol (34 µg/mL) and 0.2% w/v arabinose.

Day 6

1. Mini-prep plasmid DNA and analyze for LHR and RHR insertion FseI+XbaI and AscI+BsiWI digests, respectively.
2. Confirm correctly digesting plasmids by sequencing the RHR, and select pSN150-LHR-RHR intermediate for the next step.

Day 7: Installation of DNA encoding the sgRNA

1. Digest 200 ng pSN150-LHR-RHR plasmid with I-ppoI (Promega) in 10 µL at 37 °C for 90 min, followed by heat inactivation of enzyme at 80 °C for 20 min.
2. Obtain sgRNA-encoding DNA fragment using complementary primers in a Klenow reaction (see protocol below) or purchase as synthetic DNA fragment. (*Optional*: The Klenow fragment can be agarose gel purified, if desired.)
3. Mix 20 ng each of sgRNA-encoding DNA and digested pSN150-LHR-RHR vector with 3 µL 2x Gibson Assembly master mix and incubate at 50 °C for 1 h.
4. Transform BigEasy-TSA Electrocompetent Cells with 1 µL of the Gibson reaction mixture. Transformation conditions: 1 mm cuvette, 10 µF, 600 Ohms, 1800 V.
5. Plate on LB-agar with chloramphenicol (34 µg/mL) and incubate overnight at 30 °C.

Day 8

Pick 8 colonies and grow overnight at 30 °C in liquid LB media containing chloramphenicol (34 µg/mL) and 0.2% w/v arabinose.

Day 9

1. Mini-prep plasmid DNA and analyze for sgRNA-encoding DNA insertion by I-ppoI digest.
2. Sequence digest-positive plasmids to confirm the correct sgRNA sequence using the T7 promoter or AF443 primer.
3. Confirm retention of inserted LHR and RHR regions using FseI+XbaI and AscI+BsiWI restriction digests, respectively, in selected final plasmids.

Day 10

Maxi-prep final plasmid to obtain DNA for *P. falciparum* transfections. Plasmids are quite stable during this step. However, as prudent practice, we recommend performing restriction digests as in the previous step to verify the absence of gross deletions/rearrangements.

**D. Assembling donor vectors to engineer the *3’-UTR* of *P. falciparum* loci using pSN053/054** *(Estimated time: 10 days)*

Day 1: Installation of right homologous region (RHR)

1. Digest 200 ng pSN053/054 plasmid with I-SceI in a 10 µL at 37 °C for 90 min, followed by heat inactivation of enzyme at 80 °C for 20 min.
2. PCR the RHR from gDNA using a high-fidelity polymerase and gel purify the product. Resuspend product to 20 ng/µL.
3. Mix 20 ng each of the RHR and digested vector in 3 µL 2x Gibson Assembly master mix and incubate at 50 °C for 1 h.
4. Transform BigEasy-TSA Electrocompetent Cells with 1 µL Gibson reaction mixture. Transformation conditions: 1 mm cuvette, 10 µF, 600 Ohms, 1800 V.
5. Plate on LB-agar with chloramphenicol (34 µg/mL) and incubate overnight at 30 °C.

Day 2

Pick 8 colonies and grow overnight at 30 °C in liquid LB media containing chloramphenicol (34 µg/mL) and 0.2% w/v arabinose.

Day 3

1. Mini-prep plasmid DNA and test for RHR insertion by AflII digestion to look for a shift in size of the smaller part of the plasmid (downstream of AflII).
2. Sequence inserted RHR region for correctly digesting plasmids, and select a pSN053/54-RHR intermediate plasmid for the next step.

Day 4: Installation of left homologous region (LHR)

1. Digest 200 ng pSN053/53-RHR with FseI+BsrBI, FseI+AsiSI or FseI+BsiWI (to include either a T2A sequence, FLAG or Myc/HA epitope tag, respectively) in 10 µL at 37 °C for 90 min, followed by heat inactivation of enzymes at the appropriate temperatures for this enzyme combination.
2. PCR amplify the LHR from gDNA using a high-fidelity polymerase and gel purify. Resuspend to 20 ng/µL.
3. Mix 20 ng each of LHR and digested vector in 3 µL 2x Gibson Assembly master mix,and incubate at 50 °C for 1 h.
4. Transform BigEasy-TSA Electrocompetent Cells with 1 µL of Gibson reaction mixture. Transformation conditions: 1 mm cuvette, 10 µF, 600 Ohms, 1800 V.
5. Plate on LB-agar with chloramphenicol (34 µg/mL) and incubate overnight at 30 °C.

Day 5

Pick 8 colonies and grow overnight at 30 °C in liquid LB media containing chloramphenicol (34 µg/mL) and 0.2% w/v arabinose.

Day 6

1. Mini-prep plasmid DNA and analyze for LHR insertion (NotI+BsiWI digest) and RHR retention (AflII digest-look for a shift in size downstream of Aflll).
2. Confirm correctly digesting plasmids by sequencing the LHR, and select pSN150-LHR-RHR intermediate for the next step.

Day 7: Installation of DNA encoding the sgRNA

1. Digest 200 ng pSN053/054-LHR-RHR plasmid with AfIII or I-ppoI in 10 µL at 37 °C for 90 min, followed by heat inactivation of enzyme at 80 °C for 20 min.
2. Obtain sgRNA-encoding DNA fragment using complementary primers in a Klenow reaction (see protocol below) or purchase as synthetic DNA fragment. *Optional*: The Klenow fragment can be agarose gel purified, if desired.
3. Mix 20 ng each sgRNA-encoding DNA and digested pSN150-LHR-RHR vector with 3 µL 2x Gibson Assembly master mix and incubate at 50 °C for 1 h.
4. Transform BigEasy-TSA Electrocompetent Cells using 1 µL Gibson reaction mixture. Transformation conditions: 1 mm cuvette, 10 µF, 600 Ohms, 1800 V.
5. Plate on LB-agar with chloramphenicol (34 µg/mL) and incubate overnight at 30 °C.

Day 8

Pick 8 colonies and grow overnight at 30 °C in liquid LB media containing chloramphenicol (34 µg/mL) and 0.2% w/v arabinose.

Day 9

1. Mini-prep plasmid DNA and analyze for sgRNA-encoding DNA insertion by AflII (or I-ppoI) digest.
2. Sequence digest-positive plasmids to verify the sgRNA sequence using T7 promoter or other user-designed primer.
3. Confirm retention of inserted LHR and RHR regions using NotI+BsiWI and AscI+I-SceI restriction digests, respectively, in selected final plasmids.

Day 10

Maxi-prep final plasmid to obtain DNA for *P. falciparum* transfections. Plasmids are quite stable during this step. However, as prudent practice, we recommend performing restriction digests as in the previous step to verify the absence of gross deletions/rearrangements.

**E. Assembling donor vectors to achieve dual TetR aptamer-mediated target gene regulation at their native loci using pSN053/054** *(Estimated time: 13 days)*

Day 1: Installation of right homologous region (RHR)

1. Digest 200 ng pSN053/054 plasmid with I-SceI in 10 µL at 37 °C for 90 min, followed by heat inactivation of enzyme at 80 °C for 20 min.
2. PCR the RHR from gDNA using a high-fidelity polymerase and gel purify. Resuspend to 20 ng/µL.
3. Mix 20 ng each of RHR and digested vector with 3 µL 2x Gibson Assembly master mix and incubate at 50 °C for 1 h.
4. Transform BigEasy-TSA Electrocompetent Cells with 1 µL Gibson reaction mixture. Transformation conditions: 1 mm cuvette, 10 µF, 600 Ohms, 1800 V.
5. Plate on LB-agar with chloramphenicol (34 µg/mL) and incubate overnight at 30 °C.

Day 2

Pick 8 colonies and grow overnight at 30 °C in liquid LB media containing chloramphenicol (34 µg/mL) and 0.2% w/v arabinose.

Day 3

1. Mini-prep plasmid DNA and test for RHR insertion by AflII restriction digestion.
2. Sequence inserted RHR region of correctly digesting plasmids, and select a pSN053/54-RHR intermediate plasmid for the next step.

Day 4: Installation of left homologous region (LHR)

1. Digest 200 ng pSN053/054-RHR with FseI in 10 µL at 37 °C for 90 min, followed by heat inactivation of enzyme at 80 °C for 20 min.
2. PCR amplify LHR from gDNA using a high-fidelity polymerase and gel purify. Resuspend to 20 ng/µL.
3. Mix 20 ng each of LHR and digested vector in 3 µL 2x Gibson Assembly master mix, and incubate at 50 °C for 1 h.
4. Transform BigEasy-TSA Electrocompetent Cells with 1 µL Gibson reaction mixture. Transformation conditions: 1 mm cuvette, 10 µF, 600 Ohms, 1800 V.
5. Plate on LB-agar with chloramphenicol (34 µg/mL) and incubate overnight at 30 °C.

Day 5

Pick 8 colonies and grow overnight at 30 °C in liquid LB media containing chloramphenicol (34 µg/mL) and 0.2% w/v arabinose.

Day 6

1. Mini-prep plasmid DNA and analyze colonies for LHR insertion NotI+FseI digest and RHR retention AflII digest.
2. Confirm LHR sequence for correctly digesting plasmids, and select pSN150-LHR-RHR intermediate for the next step.

Day 7: Installation of recoded DNA encoding CDS for gene of interest

1. Digest 200 ng pSN053/054-LHR-RHR vector with BsrBI/AsiSI/BsiWI in 10 µL at 37 °C for 90 min, followed by heat inactivation of enzyme(s) at the appropriate temperature for 20 min. See below the epitope tags that can be used based on the restriction enzyme used.

| Restriction enzymes | Tag on N-terminus | Tag on C-terminus |
| --- | --- | --- |
| BsrBI | none | FLAG, c-Myc or HA |
| AsiSI | none | FLAG, c-Myc or HA |
| BsiWI | ± FLAG | c-Myc or HA |

2. Resuspend the synthetic gene to 20 ng/µL in water. Note: ensure the synthetic gene has Gibson homology to the left and right Gibson regions.

3. Mix 20 ng each of synthetic gene DNA and digested vector with 3 µL 2x Gibson Assembly master mix and incubate at 50 °C for 1 h.

4. Transform BigEasy-TSA Electrocompetent Cells with 1 µL of Gibson reaction mixture. Transformation conditions: 1 mm cuvette, 10 µF, 600 Ohms, 1800 V.

5. Plate on LB-agar with chloramphenicol (34 µg/mL) and incubate overnight at 30 °C.

Day 8

Pick 8 colonies and grow overnight at 30 °C in liquid LB media containing chloramphenicol (34 µg/mL) and 0.2% w/v arabinose.

Day 9

1. Mini-prep plasmid DNA and analyze colonies for insertion of recodonized CDS by BsiWI and NotI digestion. Use the pSN053/054-LHR-RHR plasmid as a control to detect increase in size of diagnostic fragment containing the inserted CDS.
2. Sequence positive clones to verify correct CDS sequence.
3. Confirm retention of LHR and RHR regions using NotI+BsrBI and AflII restriction digests, respectively.
4. Select pSN053/054-LHR-rCDS-RHR intermediate plasmid for the next step.

Day 10: Installation of DNA encoding the sgRNA

1. Digest 200 ng pSN053/054-LHR-rCDS-RHR plasmid with AfIII or I-ppoI in 10 µL at 37 °C for 90 min, followed by heat inactivation of enzyme at 80 °C for 20 min.
2. Obtain sgRNA-encoding DNA fragment using complementary primers in a Klenow reaction (see protocol below) or purchase as synthetic DNA fragment. *Optional*: The Klenow fragment can be agarose gel purified, if desired.
3. Mix 20 ng each sgRNA-encoding DNA and digested pSN053/054-LHR-rCDS-RHR vector with 3 µL 2x Gibson Assembly master mix and incubate at 50 °C for 1 h.
4. Transform BigEasy-TSA Electrocompetent Cells with 1 µL of the Gibson reaction mixture. Transformation conditions: 1 mm cuvette, 10 µF, 600 Ohms, 1800 V.
5. Plate on LB-agar with chloramphenicol (34 µg/mL) and incubate overnight at 30 °C.

Day 11

Pick 8 colonies and grow overnight at 30 °C in liquid LB media containing chloramphenicol (34 µg/mL) and 0.2% w/v arabinose.

Day 12

1. Mini-prep plasmid DNA and analyze colonies for sgRNA-encoding DNA insertion by AflII (or I-ppoI) digestion.
2. Sequence digest-positive clones to verify correct sgRNA identity using T7 promoter or other user-designed primer.
3. Confirm retention of inserted LHR, rCDS and RHR regions using NotI+BsrBI, BsrBI+XmaI and I-CeuI restriction digests, respectively, in the selected final pSN053-LHR-rCDS-RHR-sgRNA/pSN054-LHR-rCDS-sgRNA-RHR plasmids.

Day 13

Maxi-prep final plasmids to obtain DNA for *P. falciparum* transfections. Plasmids are quite stable during this step. However, as prudent practice, we recommend performing restriction digests as in the previous step to verify the absence of gross deletions/rearrangements.

**F. Rescuing linear plasmids into BACS** *(Estimated time: 5 days)*

Day 1

1. Digest 200 ng linear vector construct with I-SceI+NotI at 37 °C for 90 min and inactivate restriction enzymes for 20 min at 80 °C.
2. Overnight digest 200 ng pBigBOB and pAdapter with PacI and XhoI+XbaI, respectively, in 10 µL reactions at 37 °C, followed by heat inactivation of enzymes at 80 °C for 20 min. Perform all reactions in 10ul volumes. **Note**: Since pAdapter and pBigBoB are used in all rescue reactions, larger scale preparations can be performed. Digested vectors stored at −20 °C for future experiments.

Day 2

1. Combine 1 µL each of digested linear plasmid, pAdapter and pBigBoB with 3 µL 2x Gibson Assembly Master Mix and incubate at 50 °C for 1 h.
2. Transform BAC-Optimized Replicator™ v2.0 Electrocompetent cells with 1 µL Gibson reaction.
3. Plate on LB-agar with chloramphenicol (34 µg/mL) and kanamycin (50 µg/mL) and incubate overnight at 30 °C.

Day 3

Pick 8 colonies and grow overnight at 30 °C in liquid LB media containing chloramphenicol (34 µg/mL) and kanamycin (50 µg/mL)

Day 4

1. Mini-prep BAC DNA and test for insertion of the fragment released from the linear vector during I-SceI+NotI digestion (Day1, Step 1).
2. Confirm proper overall BAC topology using AvrII, PvuI and NotI/XhoI digests.

Day 5

Maxi-prep final BAC to obtain DNA for *P. falciparum* transfections. These are quite stable during this step. However, as prudent practice, we recommend repeating restriction digests as in the previous step to verify absence of gross deletions/rearrangements.

**Supplementary Table 1.**
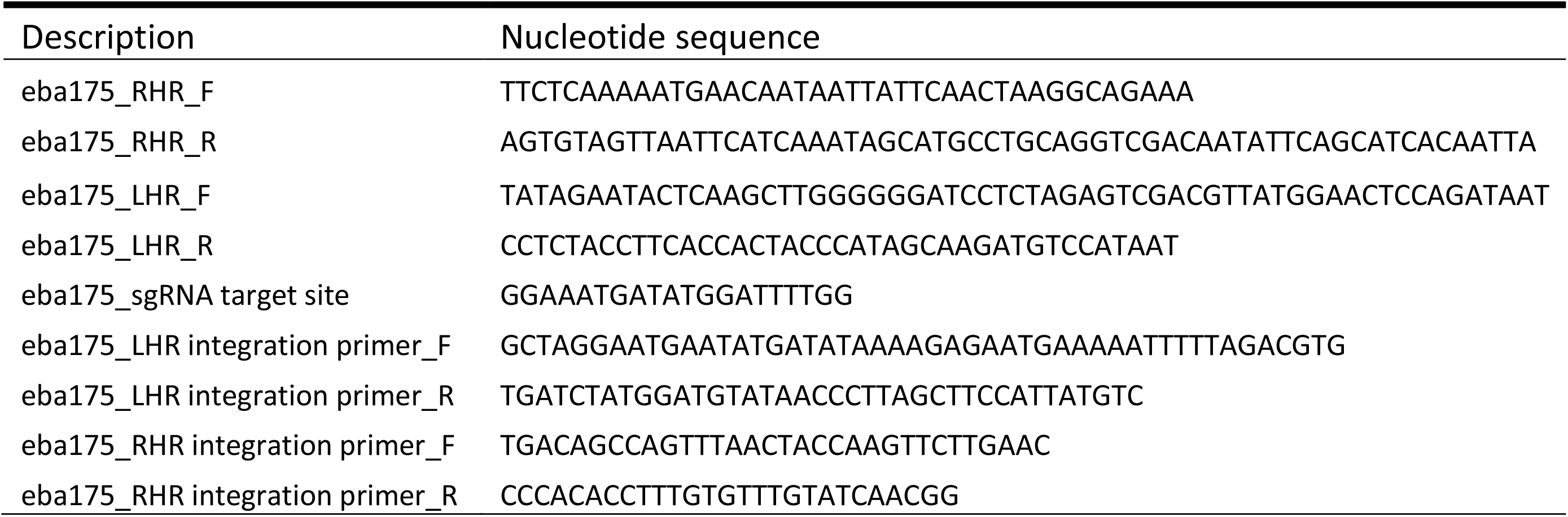
Construction of circular pUF-1 knockout vector for disrupting *eba175* locus.

**Supplementary Table 2A.**
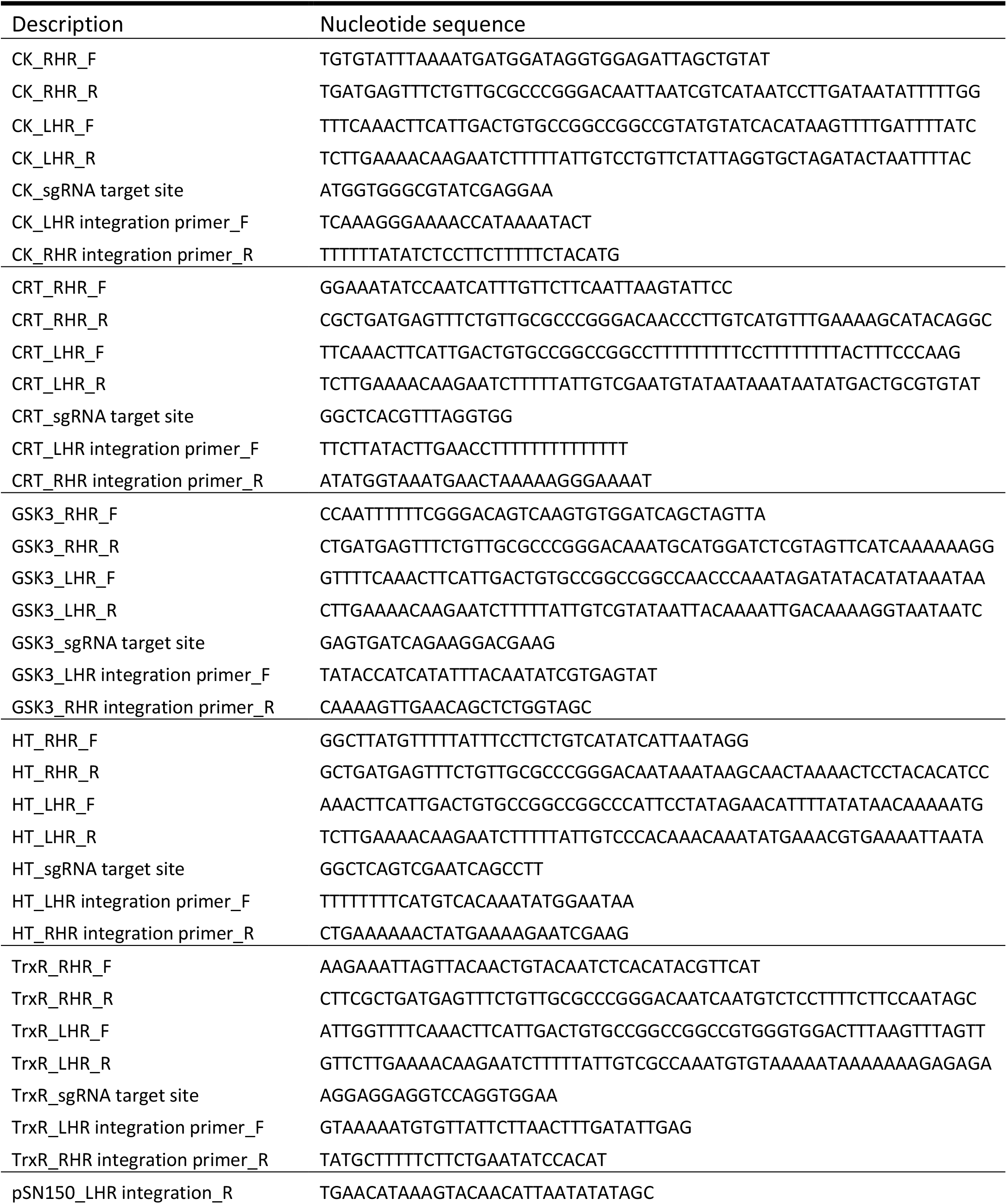
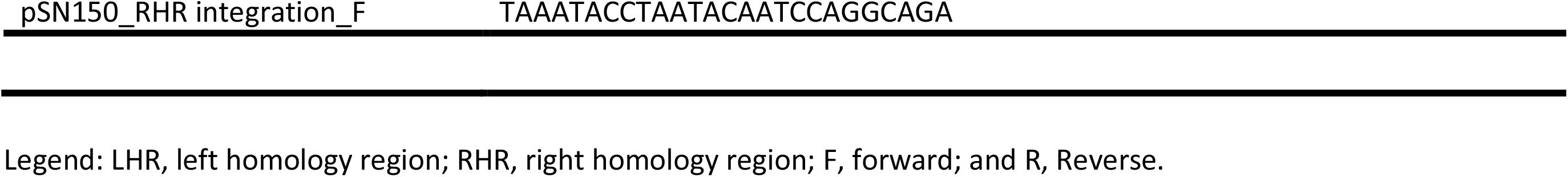
List of oligonucleotides used to construct pSN150 knockdown donor vectors and validate locus-specific integration in edited *P. falciparum* lines.

**Supplementary Table 2B.**
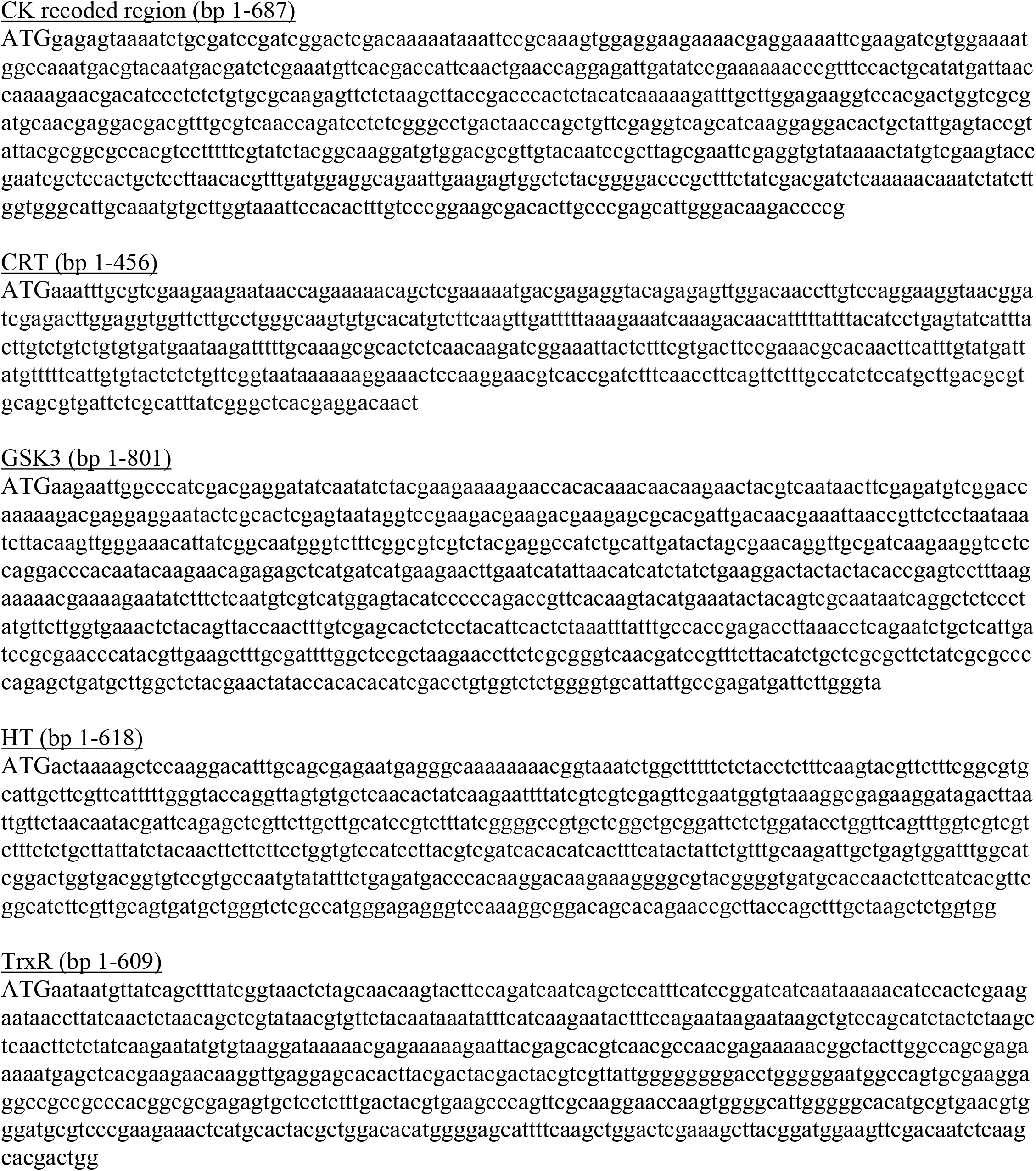
Recoded regions (*T. gondii* codon composition) used during to construct pSN150 knockdown donor vectors.

**Supplementary Table 3.**
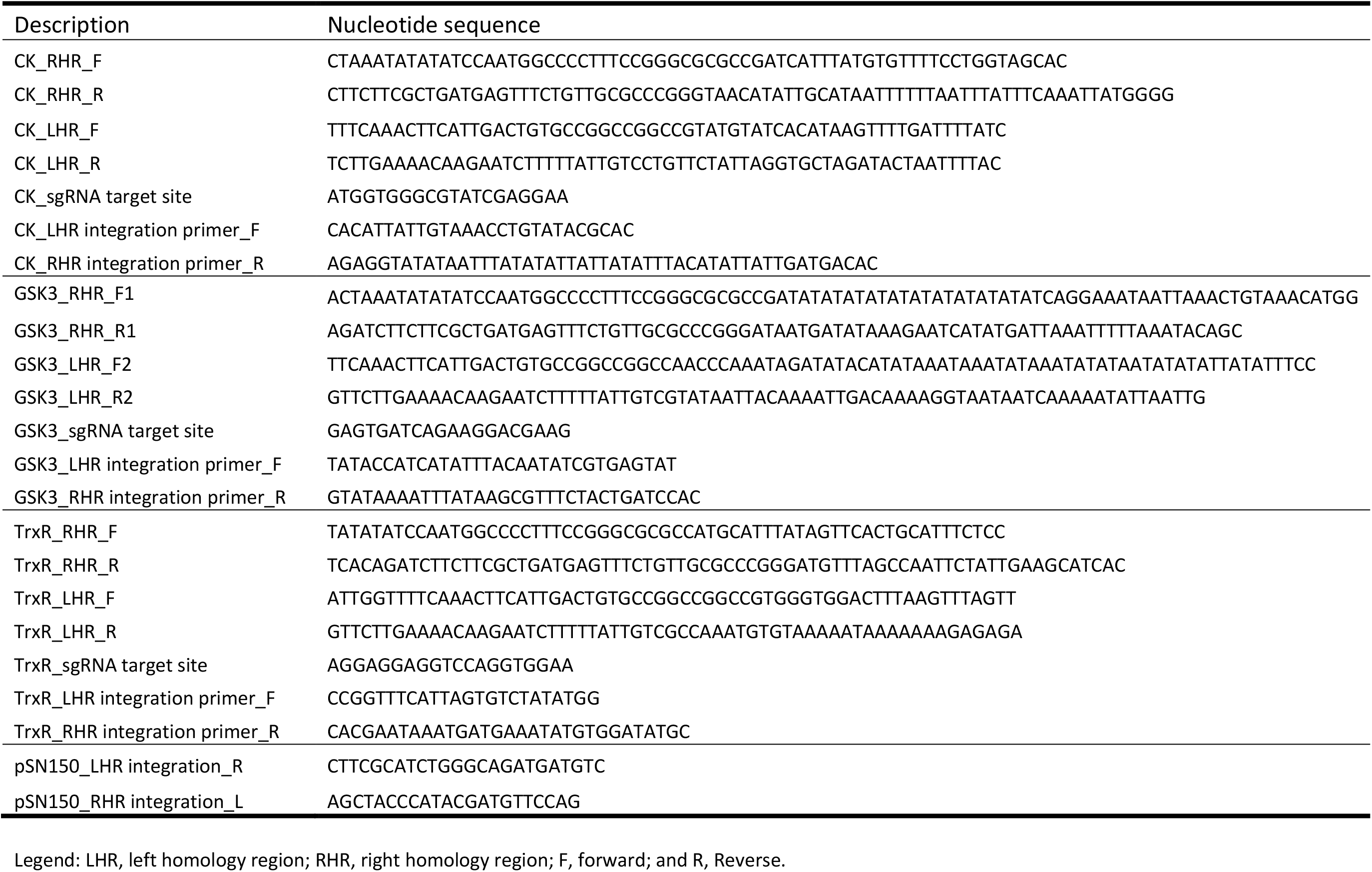
List of oligonucleotides used to construct pSN150-based knockout donor vectors and validate locus-specific integration in edited *P. falciparum* lines.

**Supplementary Table 4A.**
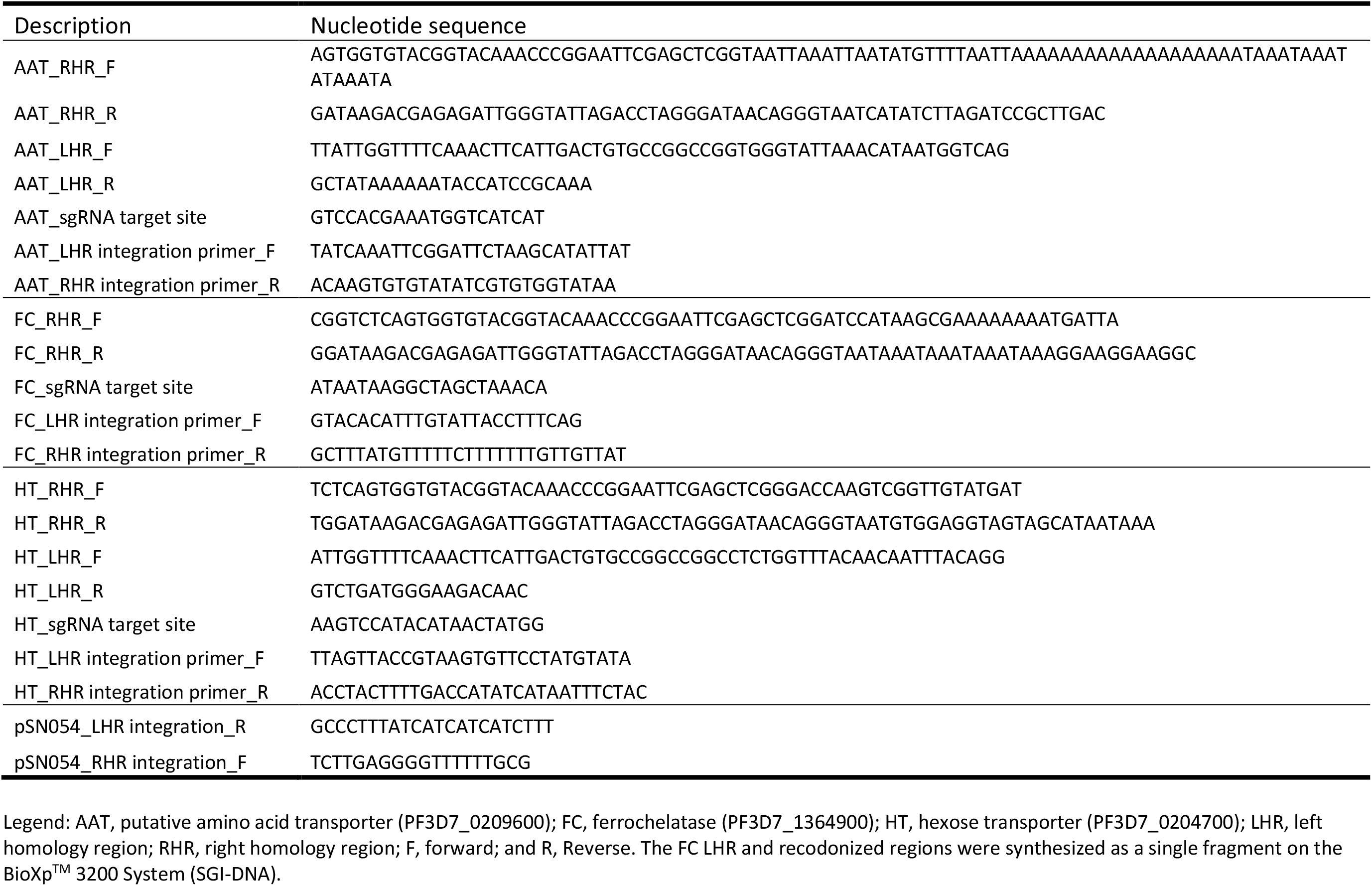
List of oligonucleotides used to construct pSN054-based knockdown donor vectors and validate locus-specific integration in edited *P. falciparum* lines.

**Supplementary Table 4B.**
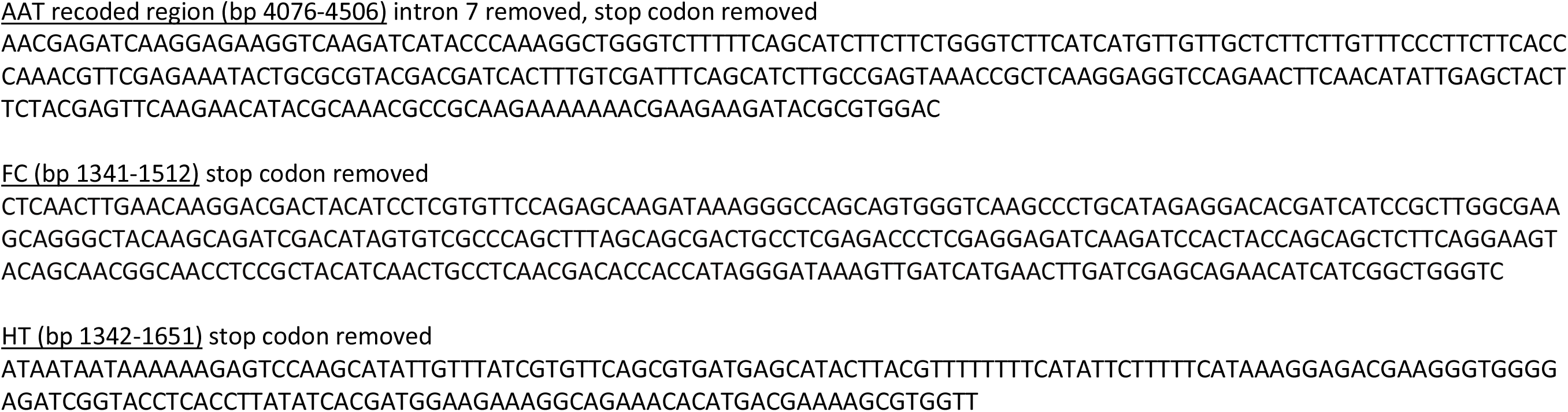
Recoded regions (*T. gondii* codon composition) used to construct pSN054-based knockdown donor vectors.

